# Alcohol Drinking Exacerbates Neural and Behavioral Pathology in the 3xTg-AD Mouse Model of Alzheimer’s Disease

**DOI:** 10.1101/726307

**Authors:** Jessica L. Hoffman, Sara Faccidomo, Michelle Kim, Seth M. Taylor, Abigail E. Agoglia, Ashley M. May, Evan N. Smith, LC Wong, Clyde W. Hodge

## Abstract

Alzheimer’s disease (AD) is a progressive neurodegenerative disorder that represents the most common cause of dementia in the United States. Although the link between alcohol use and AD has been studied, preclinical research has potential to elucidate neurobiological mechanisms that underlie this interaction. This study was designed to test the hypothesis that non-dependent alcohol drinking exacerbates the onset and magnitude of AD-like neural and behavioral pathology. We first evaluated the impact of voluntary 24-h, 2-bottle choice home-cage alcohol drinking on the prefrontal cortex and amygdala neuroproteome in C57BL/6J mice and found a striking association between alcohol drinking and AD-like pathology. Bioinformatics identified the AD-associated proteins MAPT (Tau), amyloid beta precursor protein (APP), and presenilin-1 (PSEN-1) as the main modulators of alcohol-sensitive protein networks that included AD-related proteins that regulate energy metabolism (ATP5D, HK1, AK1, PGAM1, CKB), cytoskeletal development (BASP1, CAP1, DPYSL2 [CRMP2], ALDOA, TUBA1A, CFL2, ACTG1), cellular/oxidative stress (HSPA5, HSPA8, ENO1, ENO2), and DNA regulation (PURA, YWHAZ). To address the impact of alcohol drinking on AD, studies were conducted using 3xTg-AD mice that express human MAPT, APP, and PSEN-1 transgenes and develop AD-like brain and behavioral pathology. 3xTg-AD and wildtype mice consumed alcohol or saccharin for 4 months. Behavioral tests were administered during a 1-month alcohol free period. Alcohol intake induced AD-like behavioral pathologies in 3xTg-AD mice including impaired spatial memory in the Morris Water Maze, diminished sensorimotor gating as measured by prepulse inhibition, and exacerbated conditioned fear. Multiplex immunoassay conducted on brain lysates showed that alcohol drinking upregulated primary markers of AD pathology in 3xTg-AD mice: Aβ 42/40 ratio in the lateral entorhinal and prefrontal cortex and total Tau expression in the lateral entorhinal cortex and amygdala at 1-month post alcohol exposure. Immunocytochemistry showed that alcohol use upregulated expression of pTau (Ser199/Ser202) in the hippocampus, which is consistent with late stage AD. According to the NIA-AA Research Framework, these results suggest that alcohol use is associated with Alzheimer’s pathology. Results also showed that alcohol use was associated with a general reduction in Akt/mTOR signaling via several phosphoproteins (IR, IRS1, IGF1R, PTEN, ERK, mTOR, p70S6K, RPS6) in multiple brain regions including hippocampus and entorhinal cortex. Dysregulation of Akt/mTOR phosphoproteins suggests alcohol may target this pathway in AD progression. These results suggest that nondependent alcohol drinking increases the onset and magnitude of AD-like neural and behavioral pathology in 3xTg-AD mice.

## INTRODUCTION

### Overview

The impact of alcohol use on health and well-being in older adults is not fully understood. Evidence suggests that alcohol abuse during mid-life exacerbates age-related cognitive decline and may increase the risk of developing dementia after age 65. Alzheimer’s disease (AD) is a major cause of dementia in older individuals but it is unclear if alcohol targets AD-linked molecular mechanisms to produce neural and behavioral pathology. This chapter is divided into two overall sections. First, we briefly review evidence suggesting that alcohol abuse promotes cognitive decline and dementia in older adults and present a re-analysis of our prior preclinical studies that assessed the impact of non-dependent alcohol drinking on the neuroproteome. These data show that three primary neural mechanisms of AD (Tau, amyloid precursor protein, and presenilin-1) are the statistically most likely regulators of alcohol-sensitive protein networks in the frontal cortex and amygdala of mice, which suggests that alcohol use may promote vulnerability to AD-like pathology. Second, we present results from original preclinical studies designed to evaluate the impact of non-dependent alcohol drinking on AD-like neural and behavioral pathology using the triple transgenic mouse model of AD (3xTg-AD), which expresses human Tau, APP, and PSEN-1 transgenes. This novel work suggests that alcohol use may exacerbate the onset and magnitude of AD-like pathology in vulnerable individuals. Research that focuses on the impact of alcohol use and abuse on specific molecular mechanisms of AD will move the field forward in understanding potentially unique age-dependent vulnerability in older individuals.

### Alcohol Use by Older Adults

Human beings have consumed alcohol for millennia; its use is woven into the fabric of society and religion. In 2016 approximately 43% of the global population aged 15 years and over were current alcohol drinkers with per capita intake of 32.8 grams of pure alcohol per day (WHO, 2018). Although many people drink in the absence of health consequences, alcohol misuse is associated with 5.3% of all deaths and 7.2% of premature deaths worldwide (WHO, 2018). Alcohol misuse is estimated to cost the United States $249 billion per year in health care, lost productivity, and other factors including crime and motor vehicle crashes (Sacks et al., 2015).

The negative impact of alcohol use on health depends on a variety of factors including age. Accordingly, the most recent strategic plan from the National Institute on Alcohol Abuse and Alcoholism (NIAAA; years 2017-2021) emphasizes the need for understanding the influence of alcohol on health and disease across the lifespan (NIAAA, 2017). Considerable research efforts have focused on the health effects of alcohol in prenatal, adolescent and young adult individuals. However, the percentage of the people aged 65 years or older who use alcohol is trending upward, has reached >40% of the population, and shows a positive correlation between age and drinking frequency (Breslow et al., 2017; Lewis et al., 2018). Older individuals express a variety of age-related pathologies, including neurodegenerative diseases, that increase vulnerability to other adverse health outcomes, such as poor response to and recovery from stress (Franceschi et al., 2018). Thus, alcohol use may interact with the aging process, or age-related disease conditions, to exacerbate negative health outcomes, which underscores the need for further research and public health information related to older age groups.

### Alcohol, Age-related Cognitive Decline, and Dementia

Cognitive decline is a feature of normal aging that appears to be exacerbated by alcohol use and abuse. Acute alcohol intake by older individuals is associated with reduced cognitive function and blunted neural responses that mediate higher order working memory and attention as compared to younger adults (Lewis et al., 2019; Squeglia et al., 2014). Moreover, magnetic resonance imaging studies have shown that moderate, nondependent, alcohol use (Mukamal et al., 2001) or moderate alcohol use combined with reduced physical activity (Bittner et al., 2019) is associated with loss of gray matter in older adults, which may contribute to alcohol-induced cognitive decline. However, other evidence suggests that moderate alcohol use does not alter age-related cognitive decline (Moussa et al., 2014) suggesting that the conditions under which low to moderate levels of alcohol use may impact health and well-being in older populations are not fully understood.

Alcohol abuse and dependence are associated with a higher risk of cognitive decline and dementia. For instance, 78% of adults in their upper 70’s with a history of alcohol abuse showed both cognitive impairment and increased rates of multiple forms of dementia as compared to age-matched individuals with no history of alcohol abuse (Thomas and Rockwood, 2001). In general, heavy alcohol use is associated with dementia in older populations (e.g., (Mukamal et al., 2001; Truelsen et al., 2002)), whereas light to moderate drinking appears to have no impact or may be protective (Ruitenberg et al., 2002).

Other data indicate that alcohol abuse in mid-life may exacerbate age-related cognitive decline and subsequent occurrence of dementia after age 65. For instance, a population-based study conducted in Australia showed that depression, bipolar disorder, anxiety, cerebrovascular disease, smoking, and alcohol dependence occurring during the mid-life decade of 55 – 65 years of age were all associated with significantly greater occurrence of dementia in people aged 65 – 69 (Zilkens et al., 2014). In that study, alcohol dependence showed an odds ratio of 4.14 (414% increase in probability) for development of dementia at age 65 that declined to 1.51 (51% increase in probability) by age 80 suggesting that the impact of alcohol on dementia risk is age-dependent both in terms of mid-life exposure and age of disease expression. Overall, evidence suggests that alcohol abuse may exacerbate age-related cognitive decline and dementia in older adults but the specific conditions under which these negative health consequences emerge and the underlying neural mechanisms remain to be fully characterized.

### Alzheimer’s Disease

Alzheimer’s disease is a progressive and irreversible neurodegenerative disorder that represents the most common cause of dementia in older populations (Alzheimer’s-Association, 2016) and is the sixth leading cause of death, accounting for 93,541 fatalities in 2014 – a 54% increase since 1999 (Taylor et al., 2017). The National Institute on Aging (NIA) estimates that over 5 million Americans have Alzheimer’s disease, but this number may triple by 2050.

Alzheimer’s disease is classified into three broad stages: 1) preclinical Alzheimer’s disease; 2) mild cognitive impairment (MCI) due to Alzheimer’s disease; and 3) dementia due to Alzheimer’s disease (Albert et al., 2011; McKhann et al., 2011; Sperling et al., 2011). Clinical symptoms are absent in the preclinical stage, but brain pathology may be present. MCI of the Alzheimer’s type involves memory disruption and difficulty in thinking. The progression of dementia due to Alzheimer’s disease ranges from mild to moderate to severe (Albert et al., 2011; McKhann et al., 2011; Sperling et al., 2011). In the mild stage of dementia, individuals suffer from impaired memory and sensory processing but can engage in normal activities with some assistance. Moderate Alzheimer’s dementia includes continued cognitive decline, impaired short- and long-term memory, loss of ability to communicate effectively and complete daily tasks. Individuals with severe dementia show worsening cognitive deficits with pronounced physical decline that includes loss of motor function and inability to eat and drink (Alzheimer’s-Association, 2019). Anxiety and fear are common symptoms of neurodegenerative disorders including Alzheimer’s disease (Chung and Cummings, 2000; Ferretti et al., 2001). Moreover, Alzheimer’s disease patients ranging from mild to severe stages exhibit apathy (72% of patients), agitation (60%), anxiety (48%), irritability (42%), aberrant motor behavior (38%), delusions (22%), and hallucinations (10%) (Mega et al., 1996) indicating that disease progression is associated with a spectrum of neuropsychiatric abnormalities.

Prominent pathophysiological characteristics of Alzheimer’s disease include plaque and neurofibrillary tangle (NFT) formation. Plaques are formed by abnormal deposits of amyloid β protein (Aβ) in specific brain regions (Benilova et al., 2012; Hardy and Selkoe, 2002). Aβ is derived by cleavage of amyloid precursor protein (APP) by β- or γ-secretase (e.g., PSEN-1). The long form of Aβ (Aβ42) is critical for plaque formation and is elevated in relation to the short Aβ40 form in the brain of AD patients. Neurofibrillary tangles consist of hyper-phosphorylated microtubule-associated protein Tau (pTau) (Ballatore et al., 2007). Numerous studies have linked the expression of plaques, tangles, and the progression of cognitive decline in Alzheimer’s disease with Aβ and Tau expression (Buckley et al., 2016).

### Alcohol and Alzheimer’s Disease: Need for Preclinical Research

Despite an abundance of research, the etiology of Alzheimer’s disease is not fully understood. Evidence suggests that 5% or fewer cases of Alzheimer’s disease are “familial,” with “sporadic” classifications accounting for 95% of AD diagnoses. Thus, research is increasingly directed toward understanding the impact of the environment, lifestyles, other disease states, and medical treatments as potential modifiable risk factors for Alzheimer’s disease. A recent meta-analysis found a variety of factors related to increased risk of Alzheimer’s disease including head injury in males, diabetes mellitus, use of conjugated equine estrogen with medroxyprogesterone acetate, current smoking, and low social engagement (Hersi et al., 2017).

Accordingly, a growing number of studies have evaluated alcohol use and abuse as potential risk factors for AD. Although a variety of primary studies suggest that light to moderate alcohol drinking may protect against AD in older adults (Weyerer et al., 2011), recent systematic reviews and meta analyses have concluded that there is disagreement in the field regarding the impact of alcohol consumption on AD, with divergent evidence showing that alcohol: 1) serves a protective role against AD; 2) increases AD risk; and 3) has no association with AD (Hersi et al., 2017; Piazza-Gardner et al., 2013). One source of disagreement may derive from inconsistency in operational definitions of alcohol drinking that include imprecise groupings of patients based on light, moderate, or heavy drinking as variables. Similarly, there is variation in measures used to diagnose AD, which rely partly on inconsistent clinical evaluations ranging from telephone interviews to neurological exam. Overall, validity and consistency of alcohol use and AD diagnostic criteria represent limitations of this research and support the need for standardized clinical and research practices (Piazza-Gardner et al., 2013).

We propose, therefore, that preclinical research strategies that utilize controlled methods of alcohol exposure and evaluate standardized mechanisms of AD pathology have exceptional potential to move the field forward in understanding the impact of alcohol use on AD-like neural and behavioral pathology.

### Rationale for the Present Study: Mechanisms of Alzheimer’s Disease Identified as Upstream Regulators of Alcohol-sensitive Protein Networks in C57BL/6J Mice

As noted above, a prominent feature of AD neural pathology is the coordinate accumulation of amyloid-β (Aβ) protein (Hardy and Selkoe, 2002) and increased expression or phosphorylation of Tau protein (Ballatore et al., 2007), which promotes plaque and NFT formation, respectively. As part of ongoing work to discover neural mechanisms of alcohol addiction, we assessed the impact of voluntary alcohol intake on the prefrontal cortex (PFC) and amygdala (AMY) proteome in C57BL/6J mice (Agoglia and Hodge, 2017; Salling et al., 2016). These brain regions were chosen for analysis based on known involvement in the rewarding, or reinforcing, effects of alcohol (Agoglia et al., 2015a; Agoglia et al., 2015b; Besheer et al., 2003; Faccidomo et al., 2016; Faccidomo et al., 2015a; Hodge et al., 1996; Olive and Hodge, 2000; Salling et al., 2017; Schroeder et al., 2003; Schroeder et al., 2008; Ueno et al., 2001). In each study, non-dependent mice consumed alcohol (10% v/v) vs water or water only in the home-cage for ∼1 month (**Figure 1A**). Changes in the AMY or PFC neuroproteome were assessed 24-h post alcohol via two-dimensional differential in-gel electrophoresis (2D-DIGE) with protein identification by MALDI/TOF/TOF mass spectrometry (**Figure 1B**). Alcohol intake (∼12 g/kg/day) consistently (n=4 replicate gels per brain region) altered an array of fundamental protein networks that regulate a range of diseases, biological functions, molecular and cellular functions, and developmental processes suggesting that alcohol may impact health and vulnerability to disease across the lifespan (Agoglia and Hodge, 2017; Salling et al., 2016).

**Figure 1.**
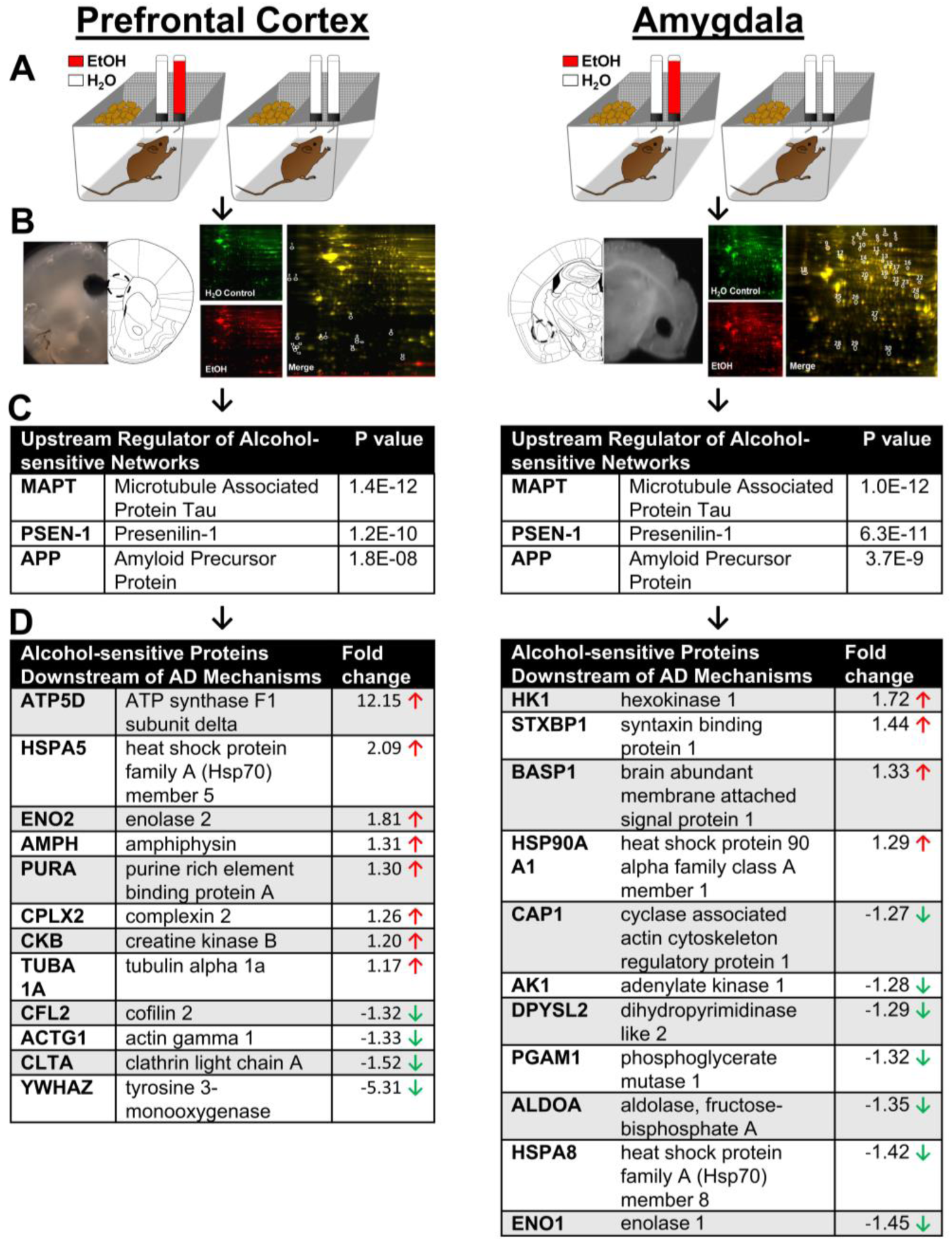
Bioinformatic analysis of alcohol-sensitive protein networks in PFC and amygdala identified major mechanisms of AD (MAPT, APP, and PSEN1) as the statistically most likely upstream regulators. (**A**) Illustration of mouse home-cage drinking method. (**B**) Illustration of sample brain regions showing anatomical location and sample tissue punches, and representative 2D-DIGE gels from each experiment. (**C**) Results of IPA Upstream Regulator Analysis that identified Alzheimer’s disease mechanisms as top regulators of alcohol-sensitive protein networks. (**D**) Alcohol-sensitive proteins detected in each brain region that are known to be downstream of MAPT, APP, and PSEN1. P values indicate significant overlap between AD regulator protein and alcohol-sensitive Alzheimer’s-linked molecules. Arrows indicate increase (↑) or decrease (↓) in expression after alcohol drinking. Data are shown as fold change representing the mean of 4 replicate gels. Proteins were included only if significantly changed in all 4 gels.

Here, we reanalyzed fold-change data (alcohol intake versus control) from the alcohol-sensitive protein networks (Agoglia and Hodge, 2017; Salling et al., 2016) using Ingenuity Pathway Analysis (IPA) Upstream Regulator Analysis (URA) to elucidate potential upstream regulatory molecules that may explain observed changes in protein networks (Kramer et al., 2014). In the PFC and AMY datasets, the URA identified MAPT (Tau), APP (amyloid precursor protein), and PSEN-1 (presenilin-1) as the most probable upstream modulators of alcohol-sensitive protein networks (**Figure 1C**). In the PFC, 12 Alzheimer’s-associated proteins were identified as downstream targets of Tau, APP, and PSEN-1 (**Figure 1D, left**). Similarly, 11 Alzheimer’s-associated proteins were identified in the AMY as downstream targets of Tau, APP, and PSEN-1 (**Figure 1D, right**). These findings indicate that alcohol intake influences protein networks that are downstream of primary molecular determinants of AD, which suggests that alcohol may influence risk of AD-related pathology in the PFC and AMY.

In addition to their importance in Alzheimer’s disease, these proteins regulate critical molecular and cellular functions in the brain. IPA Core Analysis showed that the alcohol-sensitive protein clusters play important roles in energy metabolism and homeostasis (ATP5D, HK1, AK1, PGAM1, CKB), cytoskeletal development and function (BASP1, CAP1, DPYSL2[CRMP2], ALDOA, TUBA1A, CFL2, ACTG1), cell signaling and endocytosis (STXB1[MUNC-18], ALDOA, AMPH, CPLX2, CLTA), cellular/oxidative stress (HSPA5, HSPA8, ENO1, ENO2), transcription and DNA regulation (PURA, YWHAZ). While we did not find direct overlap between the significantly changed proteins in the mPFC and AMY, similar protein families emerged, including the heat shock proteins, proteins involved in the SNARE complex, and enolase proteins suggested related mechanisms of action between brain regions.

Predictably, many of the alcohol-sensitive proteins are abnormally expressed in the brains of AD patients. Indeed, proteomic studies of CSF/plasma from human Alzheimer’s patients (Robinson et al., 2017) and transgenic mouse models of AD (Chang et al., 2013; Fu et al., 2015; Takano et al., 2013) show considerable overlap between AD-linked protein networks and the alcohol-sensitive networks shown here. Notably, heat shock proteins and creatinine kinases were upregulated and total levels of DPYSL2 (CRMP2) a microtubule protein, was decreased in the proteomics study but is consistently hyperphosphorylated in postmortem human brains and transgenic lines. This is a consistent, yet curious, finding because heat shock proteins are thought to protect the brain from increasing amounts of oxidative stress and toxicity due to accumulated Aβ aggregates (Rivera et al., 2018). It is possible that HSP’s are increasing in a compensatory fashion in response to both the detrimental consequences of alcohol and of increasing levels of AΒ in the brain. Likewise, total levels of CRMP2 may be blunted in the alcohol-drinking mice because more of the protein may be phosphorylated and starting to aggregate with Tau and AΒ proteins. Both heat shock and CRMP proteins appear to react quickly to stress and chronic alcohol drinking and given their link to AD pathology, they may be useful early biomarkers for AD-like pathology associated with alcohol use or abuse.

Further preclinical research is needed to elucidate the impact of alcohol drinking on MAPT, PSEN-1 and APP to determine if alcohol targets these systems to increase risk of developing neural and behavioral pathologies associated with AD.

### Translational Approach: Selection of the Triple-transgenic Mouse Model of AD

Although no animal model recapitulates all aspects of AD, existing transgenic mouse lines are available that express specific aspects of the disease including amyloid plaques, NFTs, neurodegeneration and behavioral deficits that are consistent with human pathology. Examples of mouse models of AD that express human APP, PSEN-1 and Tau transgenes are listed in **Table 1** (see (Webster et al., 2014) and (Jankowsky and Zheng, 2017) for more comprehensive reviews). These and other transgenic mouse lines can be used in combination with alcohol exposure methods (e.g., voluntary drinking, dependence induction, withdrawal, etc.) in preclinical studies to model the impact of alcohol use or misuse on brain and behavioral functions in AD-vulnerable individuals. This relatively unexplored strategy could move the field forward in understanding the impact of alcohol use on health and well-being in older individuals.

**Table 1.** Examples of transgenic mouse models of AD expressing human APP, APP + PS1, Tau, and APP + PS1 + Tau.

| Mouse Model | APP | PS1 | Tau | Transgene | References |
| --- | --- | --- | --- | --- | --- |
| **PDAPP** | x |  |  | huAPP770 (Ind) minigene (cDNA + introns 6–8) | (Games et al., 1995) |
| **Tg2576** | x |  |  | huAPP695 (Swe) | (Hsiao et al., 1996) |
| **APP23** | x |  |  | huAPP751 (Swe) | (Sturchler-Pierrat et al., 1997) |
| **J20** | x |  |  | huAPP770 (Swe/Ind) minigene (cDNA + introns 6–8) | (Mucke et al., 2000) |
| **APP/PS1** | x | x | | mo/huAPP695 (Swe); Tg huPSEN1 (ΔE9) | (Jankowsky et al., 2003) |
| **APP+PS1** | x | x |  | huAPP695 (Swe); huPSEN1 (L166P) | (Holcomb et al., 1999; Radde et al., 2006) |
| **5xFAD** | x | x |  | huAPP695 (Swe/Flo/Lon); huPSEN1 (M146 L/L286 V) | (Oakley et al., 2006) |
| **Tau Tg Line 43** |  |  | x | huMAPT3R0N (wt) | (Ishihara et al., 1999) |
| **PS19** |  |  | x | huMAPT4R1N (P301S) | (Yoshiyama et al., 2007) |
| **3xTg-AD** | x | x | x | huAPP695 (Swe); MAPT4R0N (P301L); Psen1 M146V knock-in | (Oddo et al., 2003a) |

Transgenic models that express APP or PSEN-1 mutations may be utilized for identifying alcohol-mediated changes in brain and behavioral function linked specifically to amyloid pathology. As discussed above, APP is cleaved by γ-secretase (e.g., PSEN-1) to form Aβ peptides (Aβ40 and Aβ42), which contribute to amyloid plaques in AD. Transgenic mouse lines that express APP or PS1 mutations, or their combination, exhibit amyloid pathology such as elevated Aβ42/Aβ40 ratio and presence of plaques. A recent study showed that APP23/PS45 mice that express a double (APP / PSEN1) transgenic mutation exhibit increased APP, β-secretase 1, Aβ40, and Aβ42 following alcohol binge-like exposure with concomitant deficits in spatial learning and memory (Huang et al., 2018). The widely used 5XFAD mouse line express a total of five human APP and PSEN-1 transgenes, which results in rapid onset of amyloid plaques throughout the cortex and hippocampus as early as two months of age (Oakley et al., 2006) and could, therefore, be used to assess site-specific amyloid pathology associated with alcohol exposure. The 5XFAD mouse line also shows a deficit in spatial memory (Xiao et al., 2015) suggesting that cognitive decline associated with amyloid pathology can be modeled in APP / PSEN-1 transgenic mouse lines.

Neurofibrillary tangles are a prominent feature of AD pathology that are formed by aggregates of hyperphosphorylated Tau protein. Transgenic mouse lines that express human Tau mutations develop tangles and show deficits in mouse behavioral tests that model cognitive decline in AD. Evidence suggests that the majority of hippocampal- and entorhinal-based cognitive deficits (>80%) in AD patients is correlated with NFTs (Giannakopoulos et al., 2003); thus, evaluating Tau-dependent neural and behavioral deficits has strong translational value. As noted above (**Figure 2**), NFT pathology is expressed in AD in a brain region-dependent manner originating in the entorhinal cortex and progressing to the hippocampus, other areas of the cortex, and limbic structures. Mouse lines that express mutations of human Tau (e.g., 3xTg-AD mice) also show this anatomical progression of pathology and are, therefore, useful for evaluating the impact of alcohol and other potential modifiable risk factors on the development of AD-like pathology.

**Figure 2.**
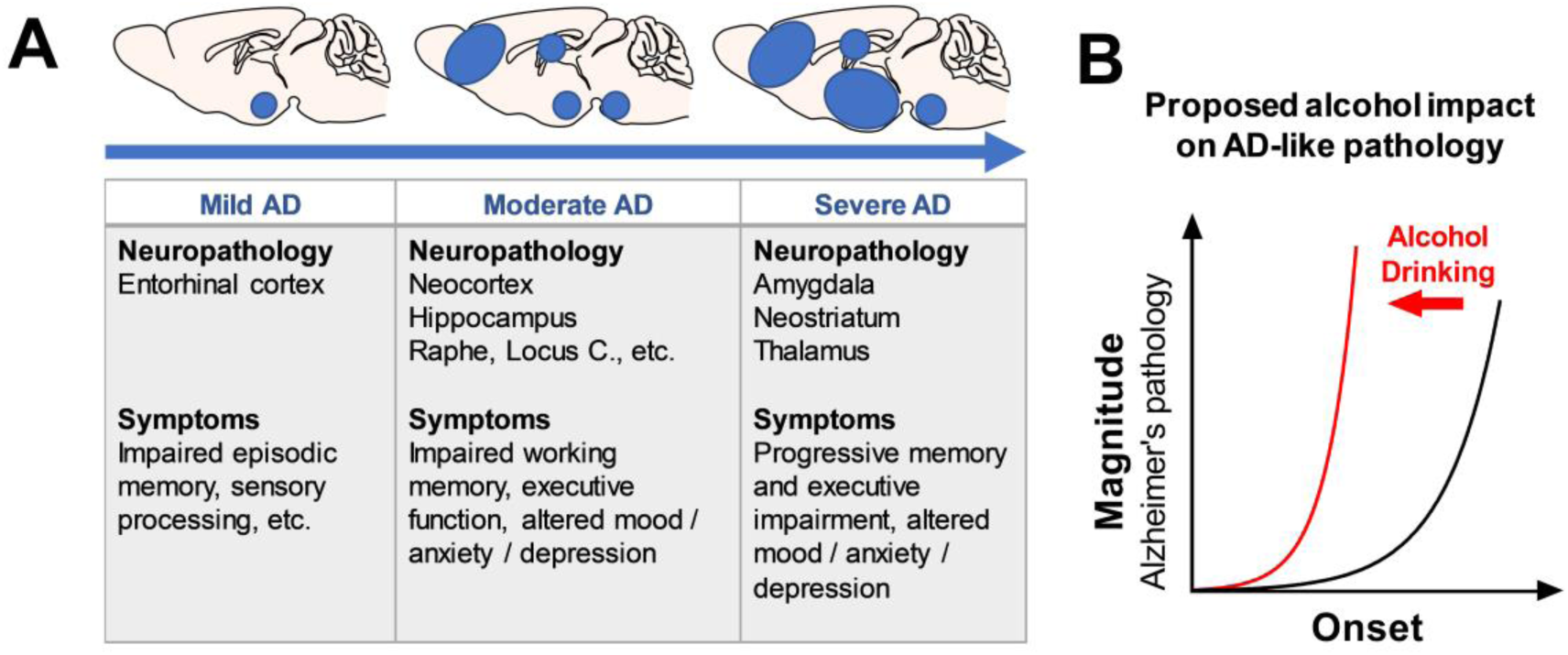
(**A**) Alzheimer’s disease progression showing mouse brain regional involvement and clinical symptoms as a function severity. (**B**) Graphic representation of the overall hypothesis of the present study: alcohol use (red line) exacerbates the onset (leftward time-dependent shift in age of onset) and magnitude (upward shift in degree) of AD-like pathology (black line).

Based on results of the URA that identified MAPT, APP, and PSEN-1 as regulators of alcohol’s impact on the neuroproteome, the present set of studies were designed to utilize the “*humanized*” triple-transgenic mouse model of AD (3xTg-AD mice). This mouse model carries a knock-in mutation of 3 human genes detected in our analysis: presenilin (PSEN-1_M146V_), amyloid beta precursor protein (APP_Swe_), and microtubule-associated protein Tau (Tau_P30IL_), expressed on a hybrid C7BL/6;129X1/SvJ;129S1/Sv genetic background (Oddo et al., 2003b). The 3xTg-AD mouse line is a well-validated animal model of AD that develops rapid age-dependent and progressive AD-like neuropathology including Aβ deposits and elevated Tau (Billings et al., 2005; Oddo et al., 2003a; Oddo et al., 2006). These neuropathies are associated with behavioral deficits including cognitive decline and altered emotional processing that are characteristic of AD (Filali et al., 2012; Pietropaolo et al., 2014; Romano et al., 2014; Webster et al., 2014). A meta-analysis of 51 studies using the 3xTg-AD mice found that Tau and Aβ (40 and 42) expression showed a strong association with impaired cognitive function (Huber et al., 2018).

Importantly, it is highly significant that 3xTg-AD mice express Tau, APP, and PSEN-1, which our neuroproteomic studies have shown are the main modulators of alcohol-sensitive protein networks within the AMY (Salling et al., 2016) and PFC of mice (**Figure 1**). To our knowledge, this innovative transgenic model of AD vulnerability has not been studied in the alcohol field leaving untapped potential for discovering alcohol-induced changes in pathology.

### Translational Approach: Analysis of Standardized Biomarkers of AD-like Pathology in 3xTg-AD Mouse Brain

As noted above, one source of disagreement in research efforts to evaluate alcohol use as a potential risk factor for AD is the lack of standardized diagnostic and research criteria (Piazza-Gardner et al., 2013). Alzheimer’s disease was originally defined as a clinical syndrome with postmortem neuropathologic verification. In 2018, the NIA and the Alzheimer’s Association (AA) jointly developed the NIA-AA Research Framework for understanding AD as an aggregate neuropathology defined by specific biomarkers in combination with postmortem examination (Jack et al., 2018). Adhering to an AT(N) rubric, it was recommended that AD research should evaluate: Aβ pathology (A); Tau pathology (T); and neurodegeneration (N) with specific biomarker combinations indicating disease state (**Table 2**).

In humans, these biomarkers can only be evaluated in plasma and CSF during life making it difficult to evaluate the influence of alcohol use on AD-related brain pathology during early disease stages. Alternatively, the present preclinical studies utilized the 3xTg-AD mouse model, to evaluate Aβ and Tau pathology in specific brain regions as a translational approach to understand the impact of alcohol drinking on disease progression. By focusing on the NIA-AA Research Framework, the present study has potential to: a) complement NIA efforts to understand causes of AD; and b) expand knowledge of alcohol-induced neuropathology in support of NIAAA efforts to understand the impact of alcohol across the lifespan.

### Overall Hypothesis: Alcohol use Impacts Alzheimer’s Disease Progression

The progression of Alzheimer’s disease is divided into stages based on the severity of behavioral and neural pathology (Braak and Braak, 1998; Palmer, 2002). AD-related neuropathies result in cell damage and death in a brain-regional progression moving from the entorhinal cortex to the hippocampus (HPC) and frontal cortex, and finally to limbic systems, including the AMY with corresponding escalation of behavioral pathology (Pietrzak et al., 2015; Sperling et al., 2011; Villemagne et al., 2013) (**Figure 2A**). Cognitive decline is initially associated with Aβ and Tau pathology in the entorhinal cortex (**mild**). With disease progression, pathology spreads to the HPC and neocortex (**moderate**) and symptoms include more prominent memory loss. Finally, in the latter stages of AD, pathology progresses to the AMY and striatum (**severe**) with increasing pathology and severity of symptoms (Palmer, 2011 #50) (**Figure 2A**).

The impact of alcohol use on Aβ and Tau pathology in specific brain regions and association with and cognitive function is not fully understood. **To address this gap in knowledge, we conducted a set of translational studies in 3xTg-AD mice designed to test the overall hypothesis that voluntary alcohol drinking exacerbates the onset and magnitude of AD-related neural and behavioral pathology (Figure 2B)**. These studies can provide significant public health information for approximately 80% of adults who have consumed alcohol in a non-dependent manner. Moreover, understanding the impact of alcohol drinking on AD pathology has potential to lead to novel diagnostic, prevention, and treatment options for this large segment of the population.

## MATERIALS & METHODS

### Mice

Male (n=7) and female (n=3) 3xTg-AD (B6;129-Tg(APPSwe, TauP301L)1Lfa *PSEN-1^tm1Mpm^*/Mmjax) triple transgenic homozygous mice were used in these studies. The 3xTg-AD mouse line was generated by Dr. Frank LaFerla at UC Irvine (Oddo et al., 2003b) and provided for use in these studies via the Mutant Mouse Resource & Research Center Repository (MMRRC stock #34830) at Jackson Labs (Bar Harbor, ME) under an approved Material Transfer Agreement with UC Irvine. Background strain- and sex-matched B6129SF2/J mice (male [n=7] and female [n=3] were used as wildtype (WT) controls where appropriate. Data from male and female mice were combined but due to low sample size, methods were underpowered to detect sex differences.

The 3xTg-AD mice are a well-validated “humanized” animal model that uniquely expresses plaques, tangles, and associated cellular and brain regional pathologies associated with Alzheimer’s disease (Oddo et al., 2003b). The 3xTg-AD mice were derived single-cell embryos from mice bearing a knockin mutation of human presenilin (PS1_M146V_) on a mixed C7BL/6;129X1/SvJ;129S1/Sv genetic background (B6;129-*PSEN-1^tm1Mpm^*). These mice were co-injected with two mutant human transgenes for amyloid beta precursor protein (APP_Swe_) and microtubule-associated protein Tau (Tau_P30IL_). Both transgenes integrated at the same locus.

All mice were at 8-weeks of age upon arrival from Jackson Labs. Mice were group-housed (n=4/cage) in techniplast cages (12” x 6.5” x 7”) until 2 weeks prior to the initiation of self-administration procedures. At all times, mice had with *ad libitum* access to food (Purina Isopro 3000) and water, the temperature of the vivarium was maintained at 21 ± 1^ο^C, humidity was maintained at 40 ± 2%, and the lights were on a reverse 12L:12D cycle (lights on at 0800). All experiments were approved by the Institutional Care and Use Committee at the University of North Carolina-Chapel Hill and were conducted in accordance with the NIH Guide for the Care and Use of Laboratory Animals (NIH Publication 80-23).

### 2-Bottle Home-Cage Alcohol or Saccharin Intake

Mice were singly housed for 2 weeks prior to initiation of home-cage alcohol drinking studies. Voluntary alcohol drinking studies were conducted as previously reported in (Agoglia et al., 2016; Agoglia et al., 2015a; Besheer et al., 2006; Faccidomo et al., 2018; Hodge et al., 1999; Holstein et al., 2011; Salling et al., 2016; Stevenson et al., 2009a; Ueno et al., 2001). Briefly, mice were given continuous 24-h access to two 50mL bottles containing sweetened alcohol (25% w/v +saccharin 0.1% w/v) or water for 4-months (120 days). Parallel control groups had continuous access to 2 bottle containing saccharin (0.1% w/v) or water. Fluid consumed was measured every 48-h and the position of the bottles (left or right) was alternated to avoid a side preference. Food was available *ad libitum*. Dependent measures included volume of fluid consumed, alcohol dose (g/kg/day) and alcohol and saccharin preference. Two-bottle choice drinking ended 24-hrs prior to the first behavioral test day (open field activity). During all behavioral testing, which occurred during 1-3 weeks post-alcohol, mice had *ad libitum* access to water only (**Figure 3**). Behavioral testing occurred in the following sequence with a minimum of 1 day in between each test: Open field locomotion, Rotarod, Morris Water Maze, Prepulse Inhibition, Fear Conditioning.

**Figure 3.**
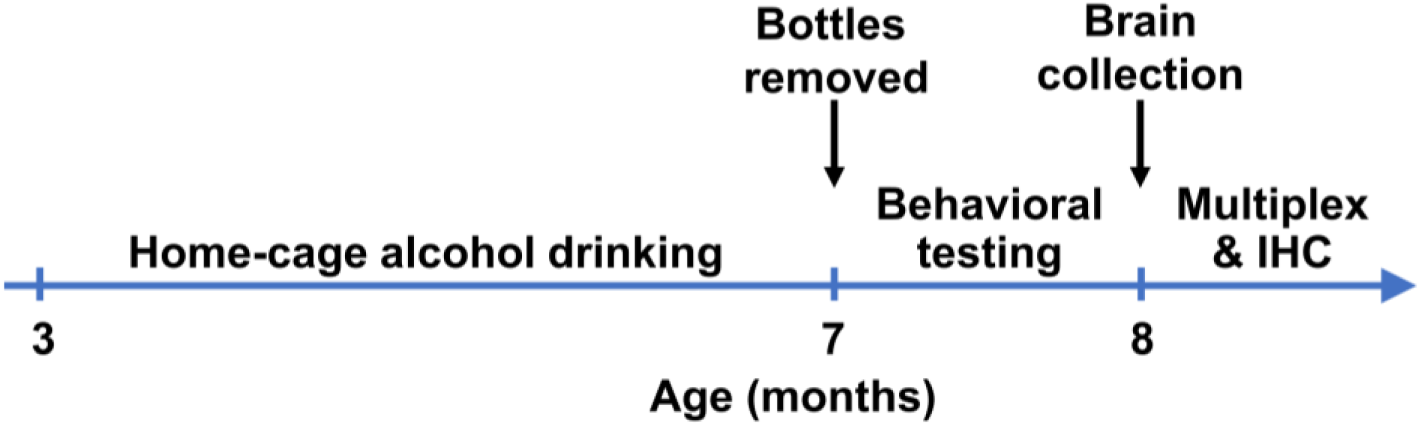
Experimental timeline. Mice drank solutions (alcohol or saccharin) starting at 3-months of age for 4-months. Behavioral testing occurred after experimental solutions were removed. Behavioral testing occurred in the following sequence with a minimum of 1 day in between each test: Open field locomotion, Rotarod, Morris Water Maze, Prepulse Inhibition, Fear Conditioning. Brains were collected at 8-months of age, 10 days after the final behavioral test

### Open Field Locomotion

Spontaneous locomotor activity and habituation to a novel environment was measured in a computer controlled open field chamber (41 cm x 41 cm x 30 cm; Versa Max, AccuScan Instruments). Locomotor distance traveled (cm) was measured by three sets of infra-red photobeams beams placed on opposite walls to record x – y ambulatory movements during a single session.

### Rotarod

Rotarod was used to assess graded motor coordination and balance in mice over 3 repeated trials. Each trial was a maximum of a 5-min test and there was a brief inter-trial interval (45-sec). Mice were placed on a rotating barrel (3 cm) of a rotarod apparatus (Ugo-Basile, Stoelting Co., Wood Dale, Il) which progressively accelerated from 3 rpm to 30 rpm. The latency to fall off or rotate around the top of the barrel was recorded as the primary measure of neuromuscular coordination.

### Morris Water Maze

This test was used to assess acquisition and memory of spatial learning. The apparatus was a large, circular pool (diameter = 122 cm; 45 cm deep), filled with water (24-26° C) and surrounded by four discrete extra-maze cues effectively dividing the apparatus into quadrants. The water was made opaque with nontoxic poster paint. A camera suspended above the maze was connected to Noldus Ethovision (Ethovision, Noldus Information Technology, Wageningen, the Netherlands). Dependent measures included swimming distance, velocity, latency to find the platform, time spent in each quadrant, path efficiency to the platform.

For *habituation and assessment of visual ability*, mice were given 4 trials in a visual platform stage. During this stage, extramaze cues were removed and the platform (12 cm in diameter) was made visible above the water surface with a flag attached to the platform. Mice were randomly placed into the pool in 1 of 4 possible locations and were given up to 60-sec per trial to find the platform. If a mouse failed to locate the platform in 60-sec, it was placed on the platform for 10-sec before being removed and allowed to rest (30-sec inter-trial interval) before starting the next trial.

For *acquisition of spatial learning*, mice were tested for the ability to find a submerged escape platform with the use of spatial cues. During this stage, extramaze cues were evenly placed around the pool to divide it into quadrants. Mice were given 4 trials daily until they reached criterion for learning (< 15-sec to find the platform). If a mouse failed to locate the platform in 60-sec, it was placed on the platform for 10s before being removed and allowed to rest (30-sec inter-trial interval) before starting the next trial. Spatial learning was demonstrated by a decrease in the latency to find the escape platform and increased efficiency in the path taken to reach the platform.

For *assessment of spatial memory*, a probe trial was conducted 1-hr after the final acquisition trial. During the probe trial, mice were placed in the pool for 60-sec in the absence of the platform but with extramaze cues in place. Time in each quadrant of the pool and the number of crossings of the target location (where the platform was located during training) was assessed and used to demonstrate memory of the platform location. Spatial memory was demonstrated by a preference for the quadrant where the platform had been located, in comparison to other quadrants of the pool.

### Fear Conditioning

This standard test was used to assess emotional processing and learning and memory. Cued fear conditioning was conducted over 3 days using an image tracking system (MED Associates, St. Albans, VT). On day 1, mice were given a 7-min training session in which they were placed in the test chamber contained in a sound attenuating box and allowed to explore for 2-min. Mice were then exposed to a 30-sec tone (80 dB), followed by a 2s foot shock (0.4 mA). Mice received an additional 2 shock-tone pairings during the training session. On day 2, mice were placed back into the test chamber and freezing behavior (immobility) was recorded as a measure of context-dependent learning across a 5-min session. On day 3, mice were evaluated for associative learning to the auditory cue across a 5-min session. Prior to the session, the chamber was modified to conceal contextual cues by using an acrylic insert to change wall and floor surfaces; additionally, a mild novel odor (dilute vanilla flavoring) was added to the sound attenuating box. Mice were then placed into the modified chamber and allowed to explore for 2-min, then the acoustic stimulus was presented for the remaining 3-min of the session. Freezing behavior before and during the stimulus were recorded.

### Acoustic Startle & Prepulse Inhibition

Prepulse inhibition was assessed as a classic measure of reflexive whole-body information processing and movement (startle response), and sensory motor gating that occurs in response to a sudden, unexpected noise. Prepulse inhibition occurs when a softer sound (the prepulse), precedes a louder sound (acoustic startle) leading to a blunted startle response in the individual.

To test this behavior in the mouse, they were placed in a small acrylic cylinder that was seated upon a piezoelectric transducer and located inside a larger sound attenuating chamber (San Diego Instruments SR-Lab system). Each chamber contained a ceiling light, fan, and a loudspeaker for the acoustic stimuli (bursts of white noise) and was connected to a computer running SR Lab software which quantifies movement and vibrations during each trial. The test session consisted of 42 trials, presented following a 5-min habituation period, with 7 different types of trials: no-stimulus trials, trials with the acoustic startle stimulus (40-ms; 120 dB) alone, and trials in which a prepulse stimulus starting 4dB above background (20-ms; 74, 78, 82, 86, 90dB) had onset 100-ms before the onset of the startle stimulus. The different trial types were presented in blocks of 7, in randomized order within each block, with an average inter-trial interval of 15-sec (range: 10 to 20sec). Measures were taken of the startle amplitude for each trial, defined as the peak response during a 65-msec sampling window that begins with the onset of the startle stimulus. Levels of percent prepulse inhibition were calculated as 100 – (prepulse response /startle response) x 100.

### Measuring NIA-AA Biomarkers in Brain

#### Multiplex array-based immunoassay of Aβ and Tau

Luminex® flow cytometry-based technology allows for simultaneous quantification of proteins in a single multiplex assay, thus providing an advantage of increased throughput and decreased volume needed per sample. This is achieved via fluorescent-coded magnetic polystyrene microspheres (80; 6.45 µm) which are coated with specific capture antibodies that allow multiple proteins to be measured via sandwich ELISAs in a single sample. Using the Luminex® 200 platform, our lab used the MILLIPLEX® MAP Human Amyloid Beta and Tau Panel (Kit # HNABTMAG-68K) to quantify Aβ40, Aβ42, Total Tau proteins (Tau), and Phosphorylated Tau Thr181 (pTau) following manufacturer protocol (Luminex® Corp, Austin TX; EMD Millipore Corporation, Billerica, MA).

Ten days after the final behavioral test (1-month post-alcohol or saccharin intake; **Figure 3**), mice (N=11) were rapidly decapitated, and brains were quickly removed and flash-frozen in cold isopentane (-40°C; 2-methylbutane; Sigma-Aldrich) for use with the multiplex assays described below. Thick coronal sections (0.8-1.0mm) were taken on a cryostat (Leica Biosystems), and using a mouse brain atlas (Franklin and Paxinos, 2008), circular tissue punches (1 mm diameter) were collected from the prefrontal cortex (PFC), medial prefrontal cortex (mPFC), nucleus accumbens (Acb), amygdala (AMY), medial hippocampus (mHPC), lateral hippocampus (lHPC), CA1 region of the hippocampus (CA1), lateral entorhinal cortex (LEC) and medial entorhinal cortex (MEC). Tissue punches from each brain region were homogenized with an ultrasonifier in 100 µl of buffer (pH 7.4) containing 20 mmol/L Tris-HCl, 7.8 mmol/L Tris-Base, 150 mmol/L NaCl, 0.02% Tween-20, and a cocktail of phosphatase and protease inhibitors (HALT) and frozen at -80°C until analysis.

Samples and reagents were thawed to room temperature and used according to manufacture protocol. Samples were diluted to 1:2 with assay buffer included in kit. Briefly, a provided lyophilized standard was reconstituted, diluted in serial fashion to create standard curves for each of the kit analytes. The standards provided quality control samples (reconstituted) and experiment samples were then added to a flat bottom 96 well plate included in the kit. Biotinylated detection antibodies were added, followed by the addition of the mixed magnetic beads. The plate was then sealed for overnight incubation at room temperature on orbital shaker (700 rpm) protected from light. The next day, the plate was washed using a handheld magnet (EMD Millipore Catalog #40-285) to retain the magnetic beads during decanting. Next, Streptavidin-PE conjugate, a reporter molecule, was added to each well and incubated on a shaker at room temperature for 30min to allow each bead to be individually identified and quantified based on fluorescent signals. After an additional round of plate washing, a final volume of wash buffer was added to the plate in place of sheath fluid for the plate to be read and discourage aggregation during the assay. The plate was read using the Luminex® 200 system equipped with xPONENT® 3.1 software. The standard curve and analyte concentrations were calculated using Milliplex Analyst 5.1 software (EMD Millipore Corporation, Billerica, MA, USA). Dilution control procedures identified a prozone effect for total Tau, which is common in Luminex and other ELISA based assays in the presence of high antigen levels (Jacobs et al., 2015), and prevented cross-plate standardization. Thus, Median Fluorescence Intensity (MFI) was measured for Tau within a single plate for each brain region analyzed.

#### pTau Immunohistochemistry (IHC)

In order to qualitatively assess levels of pTau expression in 3xTg-AD mice 1 month post alcohol- or saccharin-intake (**Figure 3**), a subset of mice (N=2) were deeply anesthetized with 100 mg/kg pentobarbital and were intracardially perfused with freshly prepared, ice cold, phosphate buffered saline (1M PBS, pH=7.4), followed by 4% paraformaldehyde (PFA). Whole brains were extracted, post-fixed in 4% PFA for 48 hrs, and then stored in PBS at 4°C. Coronal brain sections were sliced using a vibratome (Leica VT1000S) and stored in cryoprotectant (recipe) at -20°C until IHC was conducted. A subset of coronal brain sections from the anterior HPC and AMY of WT and 3xTg-AD mice with a history of saccharin or alcohol drinking were processed for fluorescent immunohistochemistry as previously published (Besheer and Hodge, 2005; Salling et al., 2016; Stevenson et al., 2009b). Briefly, sections were incubated in methanol to facilitate cell permeabilization, in hydrogen peroxide to block endogenous peroxidase activity and in a blocking buffer of Normal Goat Serum (Vector Labs) + 0.1%Triton-X in PBS (PBSTx) to inhibit non-specific binding. Next, sections were incubated overnight at RT with an anti-rabbit phosphor-Tau_Ser199,202_ polyclonal primary antibody (1:500; Invitrogen, #44-768G). The following day, sections were washed and incubated at RT with an Alexa Fluor® 488 AffiniPure Goat Anti-Rabbit IgG (1:200) secondary antibody prior to slide mounting. Fluorescence was visualized with an Olympus BX51 microscope.

### Multiplex Assay for the Akt/mTOR Signaling Pathway

To maximize efficiency and data output from our limited sample, a multiplex cell signaling assay was run on the Luminex® platform as an alternative to Western blotting. The MILLIPLEX® MAP Akt/mTOR Phosphoprotein 11-plex Magnetic Bead Kit (#48-611MAG) was used in conjunction with MILLIPLEX® MAP Phospho ERK/MAPK 1/2 (Thr185/Tyr187) Magnetic Bead MAPmate™ and MILLIPLEX® MAP Total β-Tubulin Magnetic Bead MAPmate to simultaneously detect 12 phosphoproteins [phosphorylated p70S6K (Thr412), IRS1 (Ser636), GSK3α (Ser21), GSK3β (Ser9), Akt (Ser473), PTEN (Ser380), IR (Tyr1162/Tyr1163), IGF1R (Tyr1135/Tyr1136), RPS6 (Ser235/Ser236), TSC2 (Ser939), and mTOR (Ser2448), ERK/MAPK 1/2 (Thr185/Tyr187)] with β–Tubulin serving as a loading control. The assay was conducted following manufacturer protocol (EMD Millipore Corporation, Billerica, MA, USA).

Briefly, samples were diluted to the approximately 10 ug of total protein (working range for assay was 1 to 25 μg) using provided assay buffer. Additional analyte beads (ERK and β–Tublin) were added to the 11 premixed magnetic beads and added to each well in a 96 well plate. Samples and reconstituted control cell lysates (stimulated and unstimulated) were added to the plate and incubated with bead mixture overnight (16-20hr) at 2-8°C on a plate shaker (700 rpm) protected from light. The plate was washed using a handheld magnet (EMD Millipore Catalog #40-285) to retain the magnetic beads during decanting. Biotinylated detection antibodies were added to the plate and incubated on a shaker at room temperature, protected from light for 1-hr. Detection antibody was decanted and Streptavidin-Phycoerythrin (SAPE) was added and incubated on a shaker at room temperature, protected from light for 15-min. Next, a provided amplification buffer was added and allowed to incubate on a shaker at room temperature, protected from light for 15-min. Finally, this solution was decanted, and the beads were suspended in assay buffer before being read using the Luminex® 200 system equipped with xPONENT® 3.1 software. Median fluorescence intensity was normalized against EGF stimulated control lysate and then relative quantification was calculated against the loading control.

### Drugs

Alcohol solutions (25% w/v) were prepared by diluting 95% EtOH (Pharmco Products Inc.; Brookfield, CT) with a saccharin (0.1% w/v) solution. Saccharin (0.1 % w/v; Millipore Sigma; St. Louis, MO) was prepared in water. Pentobarbital (60 mg/mL) was dissolved in ddH2O and administered via intraperitoneal injection with a 27 Ga. needle.

### General Data Analyses

Data are presented graphically and analyzed statistically using GraphPad Prism (GraphPad Prism, Chicago, IL, USA). Data were analyzed statistically by analysis of variance (ANOVA) followed by post hoc multiple comparisons where appropriate.

### Bioinformatics

The IPA Upstream Regulator analysis was conducted to identify potential mechanisms of alcohol-induced changes in the neuroproteome. This analysis identifies a cascade of upstream regulators that can explain the observed changes in protein expression. Identification of statistically most likely regulators is based on expected effects between potential regulators and their target proteins stored in the Ingenuity® Knowledge Base. The analysis examines how many known targets of each regulator are present in the dataset and compares direction of change in experimental data to what is expected from the literature in order to predict likely regulators. Overlap p-values represent the probability of upstream regulators based on significant overlap between dataset proteins and known targets of potential regulator proteins.

## RESULTS AND DISCUSSION

### Alcohol and Saccharin Intake

To explore the potential long-term impact of non-dependent alcohol drinking on specific AD biomarkers, four groups of age-matched (3-months) 3xTg-AD and WT control mice (combined male [n=14] and female [n=6]; N=20 in total) were given a choice between: 1) Alcohol [(25% w/v)+saccharin (0.1% w/v) vs. water (n=6)]; or 2) saccharin [(0.1% w/v) vs. water (n=4)] for 11-weeks according to our standard 2-bottle choice 24-h home-cage procedure (Methods; Figure 4, A &B). There were no statistically significant differences in total alcohol or saccharin intake between 3xTg-AD and WT mice (male and female data combined). Average total alcohol intake ranged from 13.7 to 17.8 g/kg/day with preference ratios (alcohol / total fluid) ranging from 36 to 39% between genotypes (**Figure 4A**). This level of alcohol intake is below the threshold for induction of physical dependence (e.g., (Ogata et al., 1972)) and observation of mice in this study detected no signs of dependence or withdrawal.

**Figure 4.**
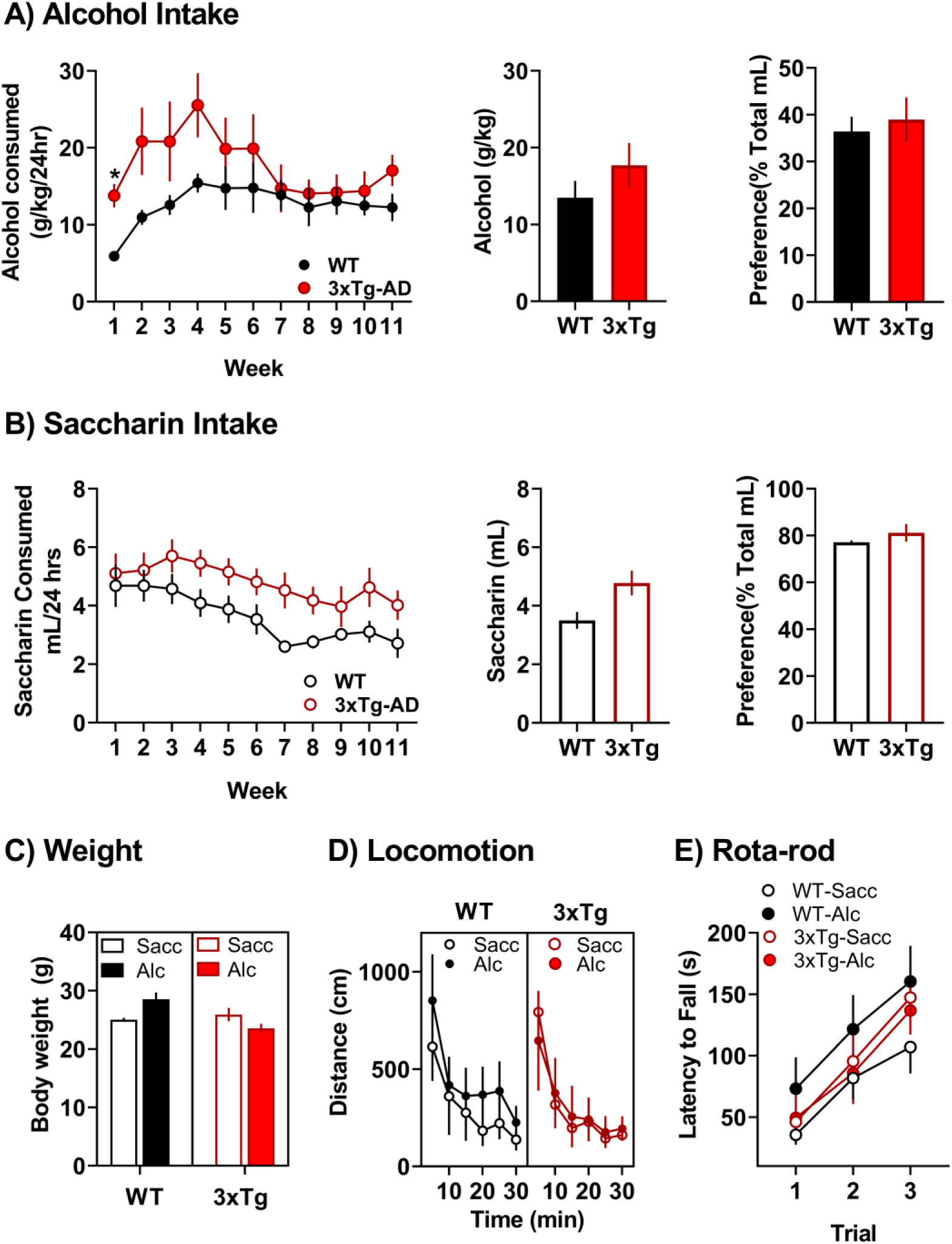
No differences in alcohol or saccharin intake or motor function in 3xTg-AD mice as compared to WT controls. (**A**) Average alcohol intake (g/kg) and preference (% Total fluid) from the 4-month access period plotted as a function of genotype. (**B**) Average saccharin intake (mL) and preference (% Total fluid) plotted as a function of genotype. (**C**) Average body weight (g) of WT and 3xTg-AD mice at 10-weeks of age (male and female combined). (**D**) Spontaneous locomotor behavior and habituation to an unfamiliar environment. Mice were placed in locomotor activity chambers, and distance traveled was measured during a single session. (E) Rotarod performance by 3xTg-AD and WT mice expressed as average latency to fall (s) over 3 trials. Data are shown as Mean ± SEM. No statistically significant differences were detected between genotypes or alcohol / saccharin condition.

Saccharin intake ranged between 3.5 to 4.8 mL/day with preference ratios between 77 and 81% between genotypes (**Figure 4B**). However, when drinking data were plotted in weekly averages (Figure 4A and B), there was a significant interaction of genotype and time (F(10,100) = 2.299). Post-hoc tests revealed that this interaction was driven by a significant difference in alcohol intake between the WT and 3xTg mice during the 1^st^ week of self-administration, with 3x-Tg mice consuming more alcohol than their wild-type counterparts. This difference was not significantly different by week 2 and by week 7, both groups consumed similar amounts of alcohol. No differences were observed in body weight between genotypes or treatment groups (**Figure 4C**). Next, during a 3-week alcohol-free period, 7-month-old mice were tested on a battery of behavioral tests in randomized order

### Normal Motor Function

Mouse models of AD show behavioral deficits on a variety of tasks that model aspects of the human disease including passive avoidance, fear conditioning, and various mazes (Webster et al., 2014). Since these and other commonly used tasks require motor function, interpretation of altered performance can be complicated by the presence of motor deficits. Evidence indicates that 3xTg-AD show reduced locomotor activity in the open field at 12 – 18 months of age (Filali et al., 2012; Gulinello et al., 2009). By contrast, 3xTg-AD mice at both 6-months (Stover et al., 2015a) and 16-months (Garvock-de Montbrun et al., 2019) of age show improved performance as compared to WT controls on the rotarod test suggesting improved motor coordination. Importantly, however, 16-month old 3xTg-AD mice show an age-dependent deficit in performance on rotarod, grip strength, gait analysis and balance beam tests as compared to younger 6-month old 3xTg-AD mice (Garvock-de Montbrun et al., 2019). Overall, these findings underscore the importance of evaluating performance of 3xTg-AD mice on AD-related cognitive tasks in the context of potential motor abnormalities.

As noted above, behavioral testing in the present study was conducted in 3xTg-AD mice and WT controls at 7 months of age. To determine if alcohol intake altered motor function in this age range, we evaluated 3xTg-AD and WT mice in the open-field and rotarod tests during the first week after alcohol exposure. In the open field test, 3xTg-AD mice showed levels of spontaneous locomotor activity and habituation that were not different than WT mice (**Figure 4D**). There was also no effect of genotype or history of alcohol intake. Three-way repeated measure (RM)-ANOVA (genotype x alcohol x time) found a significant main effect of time [F (2.040, 32.64) = 34.84, P<0.0001] with no main effect of genotype or alcohol intake, and no interactions among the variables. This indicates that mice from all groups showed similar levels of activity and habituation to the environment, which suggests the absence of deficits in gross motor function or habituation learning. When tested on the rotarod, performance of all mice improved as a function of repeated trials (**Figure 4E**). Three-way ANOVA showed a significant effect of trial [F (1.825, 29.19) = 37.74, P<0.0001], but there was no effect of alcohol intake or genotype, and no interaction among the variables. These data indicate that there are no abnormalities in motor ability among the genotypes or alcohol intake conditions at this age.

### Alcohol-induced Deficit in Spatial Memory in 3xTg-AD mice

To assess cognitive function, alcohol and saccharin exposed 3xTg-AD and WT mice were tested in the Morris Water Maze (MWM), which is a hippocampal-dependent measure of place navigation (Morris et al., 1982). The MWM is widely used as a preclinical measurement of reference memory and working memory (Webster et al., 2014). At the conceptual level, the MWM task recapitulates symptoms seen in human AD patients including confusion and getting lost due to difficulty with navigation. Numerous mouse genetic models of AD have been tested in the MWM and most show diminished performance that is consistent with AD-like cognitive deficits (e.g., (Lalonde et al., 2005; Ohno et al., 2007; Webster et al., 2014).

#### Spatial Learning

Results showed that 3xTg-AD mice exhibited a deficit in acquisition of spatial learning that was not altered by alcohol exposure (**Figure 5A, left**), which is consistent with known performance of this mouse strain (Huber et al., 2018). Three-way RM-ANOVA (genotype x alcohol x trial) identified a significant main effect of trial [F (2.006, 24.07) = 3.511, P=0.0458] and genotype [F (1, 12) = 7.841, P=0.02]. However, there was no main effect of alcohol exposure or an interaction among the variables.

**Figure 5.**
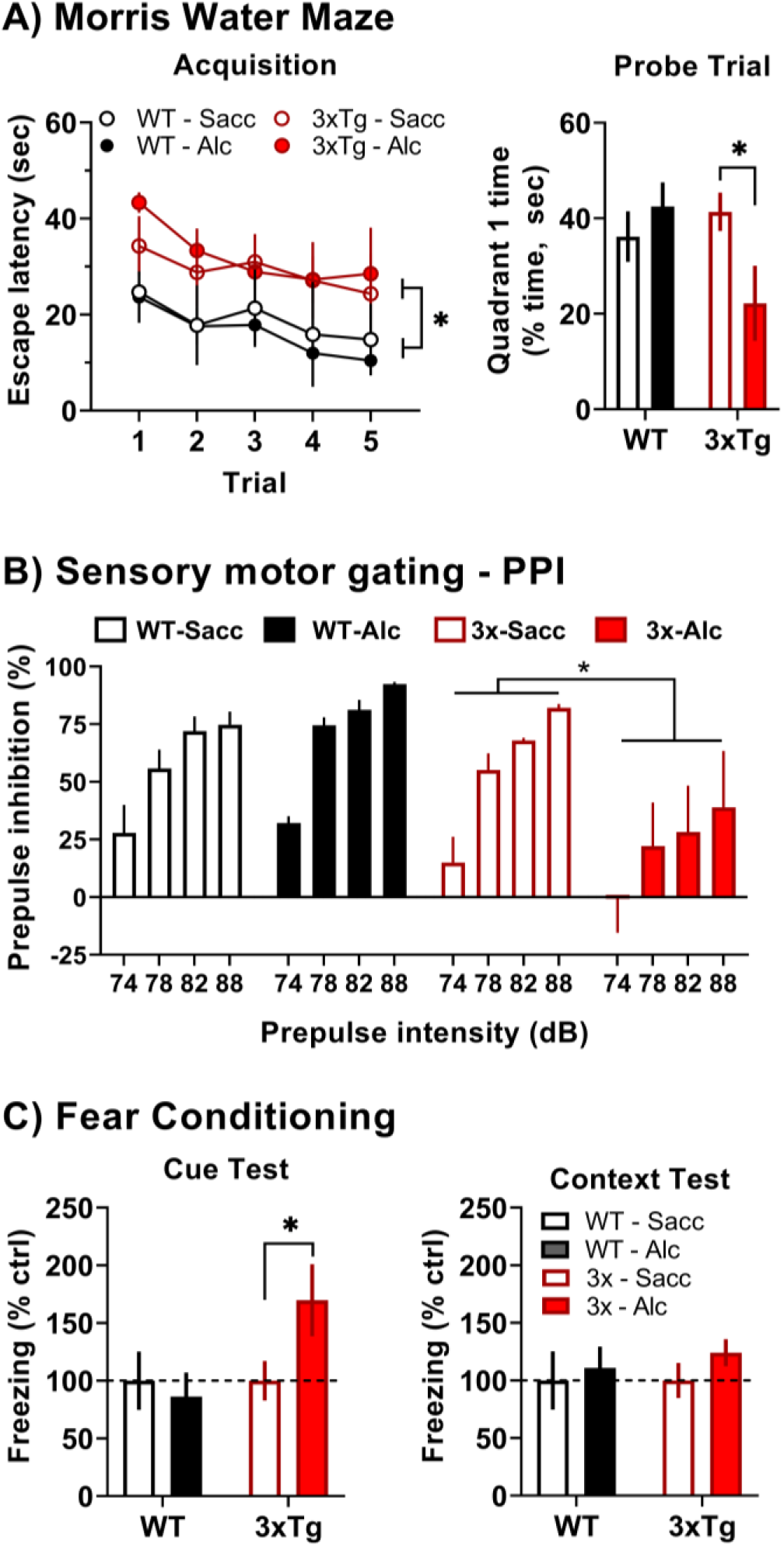
Alcohol drinking-induced behavioral deficits in 3xTg-AD mice. (**A, left**) Average escape latency (sec) plotted as a function of trial during Morris Water Maze acquisition. * indicates significant main effect between genotypes over all trials. (**A, right**) Percentage of time (sec) spent in the quadrant of the Morris Water Maze that previously contained the escape platform. * indicates significant difference between alcohol versus saccharin exposed 3xTg-AD mice, P<0.05. (**B**) Average prepulse inhibition (%) plotted as a function of stimulus intensity (dB). * indicates statistically significant main effect between 3xTg-AD mice that consumed alcohol versus saccharin. (**C, left**) Cued fear response plotted as average freezing (% control seconds) as a function of genotype and alcohol intake condition. * indicates significant difference between 3xTg-AD mice that consumed alcohol versus saccharin, P<0.05. (**C, right**) Context-dependent fear response plotted as freezing (% control) as a function of genotype and treatment conditions. Abbreviations: Saccharin (Sacc), Alcohol (Alc).

These data show that 3xTg-AD mice at 7-months of age exhibit a deficit in spatial learning as demonstrated by increased escape latencies during the acquisition phase of the MWM test irrespective of treatment condition. This result is consistent with prior evidence showing that at 6-months of age, 3xTg-AD mice require more trials to reach criterion (escape latency) when learning the MWM task, which was demonstrated by single-trial analysis to be a function of impaired memory function during the multi-day procedure (Billings et al., 2005).

Importantly, there is no evidence in the present study that a history of alcohol intake altered spatial learning by 3xTg-AD mice or WT controls. However, since neural (Oddo et al., 2003b) and behavioral (Garvock-de Montbrun et al., 2019) pathology continues to develop in 3xTg-AD mice after 12 months of age, it will be important for future studies to evaluate the impact of alcohol use on spatial learning in older mice.

#### Spatial Memory Probe Trial

When tested 1-hr after the last learning trial, alcohol-exposed 3xTg-AD mice exhibited a significant memory deficit as shown by less time spent in the quadrant previously containing the escape platform (**Figure 5A, right**), which is consistent with an alcohol-induced deficit in hippocampal-dependent spatial memory. Two-way ANOVA (genotype x alcohol exposure) found no main effect of genotype or alcohol but identified a significant interaction between those factors [F (1, 13) = 5.018, P=0.0432], indicating that the effects of alcohol depend on genotype. Multiple comparison procedures showed that alcohol exposed 3xTg-AD mice spent significantly less time in the quadrant that previously contained the escape platform (**Figure 5A, right**).

Prior research has shown that 3xTg-AD exhibit impaired spatial memory on short-term (1.5-hr) and long-term (≥ 24-hr post training) memory probe trials in the MWM (Billings et al., 2005). Our results differ from that work, however, in that only the alcohol exposed 3xTg-AD mice demonstrated a memory deficit on the probe trial conducted 1-hr post acquisition (**Figure 5A**). This deficit in performance on the probe trial could be a function of poor performance during acquisition; however, this interpretation does not seem likely since saccharin exposed (control) 3xTg-AD mice showed a similar pattern of performance during acquisition but exhibited normal performance on the memory probe trial. This suggests that the level of performance by both groups of 3xTg-AD mice during acquisition was sufficient to support normal performance on the probe trial. Overall, these data indicate that non-dependent alcohol use produced a deficit in hippocampal-dependent spatial memory.

Since deficits in spatial navigation are pronounced in AD, this finding is important from a translational perspective. The HPC is an initial locus of neurodegeneration in patients affected by Alzheimer’s disease (Braak and Braak, 1991; Whitwell, 2010) with diminished navigational ability presenting as one of the first clinical symptoms (Cherrier et al., 2001; Pai and Jacobs, 2004). Evidence indicates that AD targets specific navigational skills in humans including route learning (e.g., specific orienting skills) and topographic memory or memory for landmarks (e.g., spatial memory and perception) (Lithfous et al., 2013). Thus, the present finding that 3xTg-AD mice show diminished spatial learning irrespective of alcohol exposure is consistent with initial development of AD-like pathology in this mouse model and indicative of an AD-like pathology in orienting skills. Importantly, however, our finding that the prior alcohol exposure impaired performance on the MWM memory probe trial in 3xTg-AD mice suggests that nondependent alcohol use may specifically target spatial memory and perception in vulnerable individuals. These results agree with recent evidence from a 30-year longitudinal cohort study in humans showing that long-term moderate alcohol drinking is associated with hippocampal atrophy (Topiwala et al., 2017) and suggest that 3xTg-AD mice may show vulnerability to alcohol-induced pathology in the HPC.

### Alcohol-induced Deficit in Sensorimotor Processing

The startle reflex is an adaptive behavioral reflex that is elicited by sudden or loud stimuli. Prepulse inhibition (PPI) is a measure of sensorimotor gating where the startle reflex generated by a strong (e.g., loud) stimulus is inhibited by prior presentation of a weaker stimulus from the same sensory modality. Recent evidence indicates that patients with Alzheimer’s dementia exhibit normal acoustic startle response but show reduced PPI as compared to healthy age-matched controls or patients with mild cognitive impairment (Ueki et al., 2006). Accordingly, in the APP + PSEN-1 transgenic mouse model of AD, PPI was shown to be inversely related to Aβ expression in the cortex and HPC (Ewers et al., 2006). It is not known if there is a relationship between alcohol use and sensorimotor gating in Alzheimer’s disease.

To address this question, alcohol and saccharin exposed 3xTg-AD and WT controls were tested for PPI of the startle response. Results showed that saccharin exposed 3xTg-AD mice exhibited normal PPI, but a history of alcohol drinking was associated with a pronounced and significant impairment in PPI across a range of stimulus intensities (**Figure 5B**). Three-way RM-ANOVA (alcohol x genotype x stimulus intensity, dB) with identified a significant main effect of both genotype [F (1, 12) = 6.777, P=0.023] and dB [F (3, 36) = 44.7, P<0.0001], and a significant alcohol x dB interaction [F (1, 13) = 4.527, P=0.038].

These data indicate that a history of alcohol drinking by 3xTg-AD mice is associated with diminished PPI of the startle response. At the age tested (7 – 8 months), saccharin exposed 3xTg-AD mice did not exhibit altered PPI. This suggests that alcohol exacerbated the onset of cognitive dysfunction in the 3xTg-AD mouse model. PPI of the startle response is regulated by interconnected forebrain circuits including the PFC, entorhinal cortex, ganglia, HPC, and AMY (Swerdlow et al., 2001). Altered sensory processing is a common feature of AD and is consistent with recent evidence of PPI deficits linked to AD pathology in the lateral entorhinal cortex (LEC) of humans and mouse models (Khan et al., 2014). Indeed, entorhinal cortex lesions are associated with reduced PPI (Goto et al., 2002). However, deficits in PPI also underlie a variety of neuropsychiatric disorders including obsessive compulsive disorder, schizophrenia, anxiety, and PTSD (Geyer and Swerdlow, 2001; Hoenig et al., 2005; Kohl et al., 2013) and may reflect diminished ability to filter or process relevant environmental stimuli. Thus, the alcohol-induced reduction in PPI may reflect the impact of extended alcohol drinking across a continuum of affective and neurological disorders in 3xTg-AD mice.

### Alcohol-induced Dysregulation of Emotional Processing in 3xTg-AD Mice

Several studies have assessed cued and contextual fear conditioning in 6-month old 3xTg-AD mice as measures of emotional and cognitive processing; however, results are inconsistent. One study reported increased freezing behavior by 3xTg-AD mice when exposed to a context previously paired with shock (Espana et al., 2010), which suggests intact associative memory with enhanced fear response. By contrast, another study reported reduced latency by 3xTg-AD mice to voluntarily enter a context previously paired with shock (Billings et al., 2005), which is consistent with reduced fear or emotional response and may reflect a hippocampal-dependent deficit in spatial memory. Alternatively, some studies have reported no difference between 3xTg-AD mice and WT controls on measures of cued or contextual fear conditioning (Chu et al., 2012; Stover et al., 2015b). In general, however, both the APP and 3xTg-AD mouse models of AD show associative memory deficits and emotional dysfunction (Billings et al., 2005; Gimenez-Llort et al., 2007; Saura et al., 2005) that are associated with Aβ pathology.

In the present study, cued and contextual fear conditioning were tested in 3xTg-AD and WT mice to assess the potential impact of a history of alcohol intake on emotional responding and associative memory. Results showed heightened freezing behavior by alcohol exposed 3xTg-AD mice as compared to saccharin controls when presented with a cue that was previously paired with shock. No differences in freezing were observed when mice were exposed to the shock paired context (**Figure 5C**). Although, 2-way ANOVA (alcohol x genotype) found no main effect for either variable and no interaction following both the cue and context test, a single planned comparison showed that 3xTg-AD mice with a history of alcohol drinking exhibited increased freezing behavior following exposure to the cue as compared to saccharin exposed 3xTg-AD mice (t=1.947, df =6.076, P<0.05, **Figure 5C**).

Anxiety and fear are common symptoms of neurodegenerative disorders including Alzheimer’s disease (Chung and Cummings, 2000; Ferretti et al., 2001). Despite the well-known impact of alcohol on anxiety and emotional processing (Koob and Volkow, 2010), the influence of alcohol use on anxiety and fear associated with AD remains to be fully understood. The results of the present study show heightened cued-fear response by 3xTg-AD mice with a history of alcohol intake. This suggests that nondependent alcohol use may dysregulate emotional responding in individuals vulnerable to, or in the early stages of, AD.

Moreover, cue-specific fear suggests altered AMY function (Phillips and LeDoux, 1992) and is consistent with known effects of alcohol and amygdala-based fear responses (Agoglia and Herman, 2018; McCool et al., 2010). The AMY also regulates positive reinforcing effects of alcohol (Besheer et al., 2003; Besheer et al., 2012; Cannady et al., 2017; Salling et al., 2016; Schroeder et al., 2003) and cue-induced relapse to alcohol-seeking behavior (Cannady et al., 2011; Salling et al., 2017; Schroeder et al., 2008), which suggests common neuroanatomical mechanisms of AD and alcohol addiction.

### NIA-AA Biomarker Analysis in specific brain regions of 3xTg-AD mice (1-month post alcohol)

AD biomarkers including Aβ (40 and 42), Tau (total and pTau), in conjunction with imaging methods, can detect core features of AD and show promise for development of targeted therapeutics (Buckley et al., 2016; Jessen et al., 2014). As noted above, the NIA-AA Research Framework recommends a focus on these markers to enable a more accurate characterization and understanding of neural events that lead to AD-related cognitive and behavioral impairment (Jack et al., 2018). A recent meta-analysis of 51 peer-reviewed studies shows that neural expression of Aβ (40 and 42) and pTau (AT8 – ser 181 and ser 199) exhibit strong associations with cognitive decline in the well-validated 3xTg-AD mouse model of AD that expresses human MAPT, APP, and PSEN-1 transgenes (Huber et al., 2018). Taken together with our neuroproteomic data show that alcohol-sensitive protein networks in the PFC and AMY are downstream of the MAPT, APP, and PSEN-1, this provides a strong rationale for evaluating the impact of alcohol intake on NIA-AA biomarkers in 3xTg-AD mice. Here, we evaluated NIA-AA biomarkers in specific brain regions of 3xTg-AD mice as a translational approach to increase understanding of the impact of alcohol drinking on the trajectory of neural pathology in AD.

#### Lateral Entorhinal Cortex (LEC)

Alcohol intake significantly increased Aβ (42/40) ratio [t(7)=4.1,P=0.004)] and total Tau protein [t(7)=3.57,P=0.009] as compared to saccharin control (**Figure 6A**). This pathology profile is consistent with *Alzheimer’s disease and concomitant non-Alzheimer’s pathologic change* (e.g., A+T-(N)+) because elevated total Tau, in the absence of pTau, may reflect general neuropathology (**Table 2**). Since the initial (mild) stage of AD is characterized by Aβ and Tau pathology in the LEC (Braak and Braak, 1998; Khan et al., 2014; Palmer et al., 2011), these results support the interpretation that alcohol drinking enhanced the onset of AD-like neural pathology in 3xTg-AD mice.

**figure 6.**
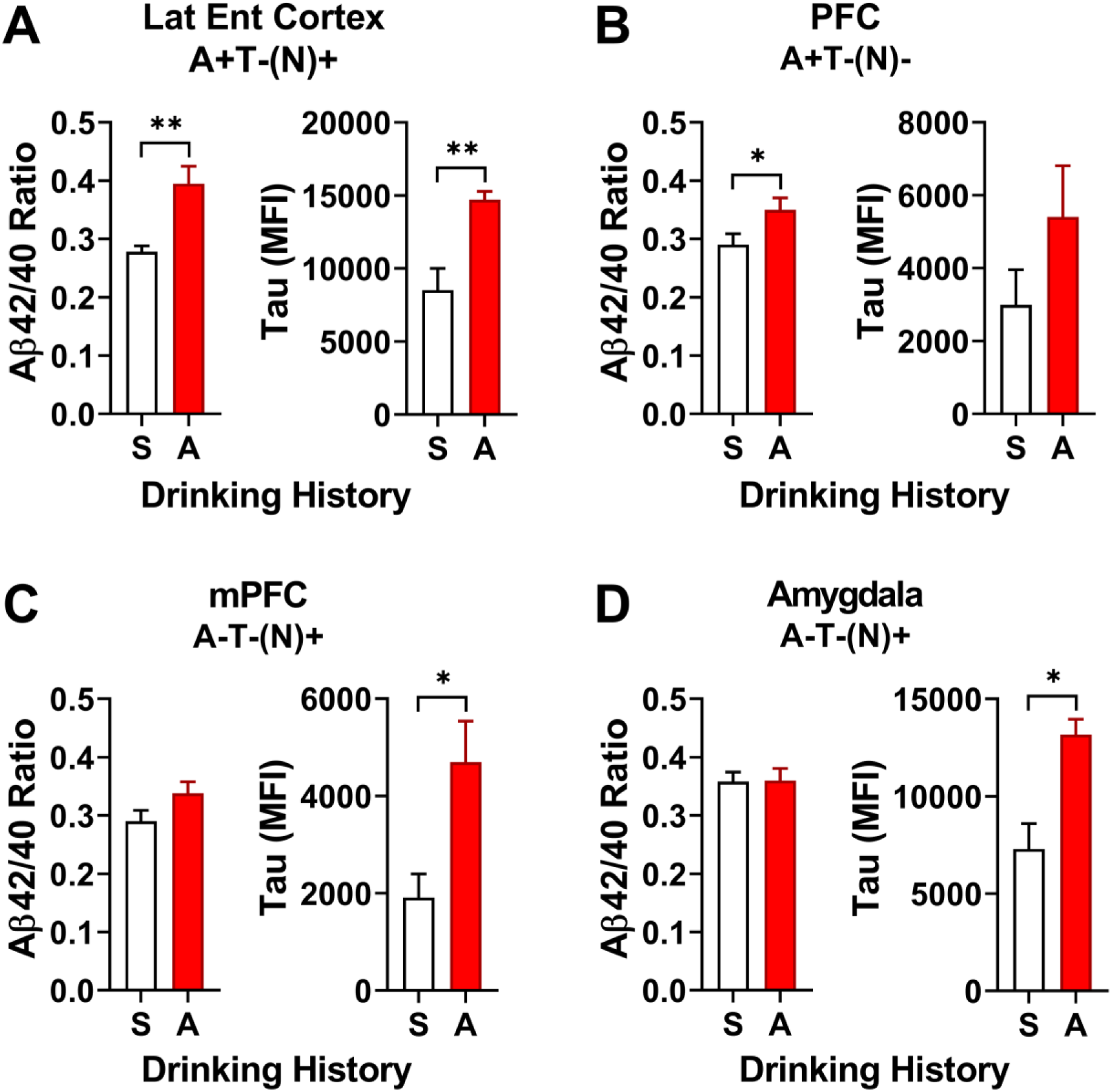
AD biomarker protein expression in specific brain regions of 3xTg-AD mice at 1-month post alcohol drinking. The title above each graph shows brain region and the AT(N) rubric for biomarker interpretation (see Table 2). (**A-D, left**) Mean ± SEM Aβ (42/40) ratio. (**A-D, right**) Total Tau protein expressed as Mean ± SEM background corrected Median Fluorescence Intensity (MFI). Measures are plotted as a function of drinking history (S – saccharin; A – Alcohol). * - P<0.05; ** - P<0.01, A vs S, t-test. WT control wells exhibited expression at background levels (data not shown).

**Table 2.** NIAA-AA Research Framework showing AT(N) biomarker groupings associated with disease categories (top) and AT(N) definitions (bottom). + indicates presence, - indicates absence of change in biomarker.

| AT(N) biomarker grouping |  | Disease category |  |
| --- | --- | --- | --- |
| A-T-(N)- |  | Normal AD biomarkers |  |
| A+T-(N)- |  | Alzheimer's pathologic change | Alzheimer's continuum |
| A+T+(N)- |  | Alzheimer's disease |  |
| A+T+(N)+ |  | Alzheimer's disease |  |
| A+T-(N)+ |  | Alzheimer's and concomitant non-AD pathology |  |
| A-T+(N)- |  | Non-AD pathology |  |
| A-T-(N)+ |  | Non-AD pathology |  |
| A-T+(N)+ |  | Non-AD pathology |  |
| AT(N) definitions |  |  |  |
| A | Aggregated Aβ (plaques); Aβ42 or Aβ42/40 ratio elevated in brain (reduced in CSF) |  |  |
| T | Aggregated Tau (tangles); elevated pTau |  |  |
| N | Neurodegeneration; elevated total Tau |  |  |
| Adapted from Jack et al., 2018. |  |  |  |

#### Prefrontal Cortex (PFC)

Alcohol intake significantly increased the Aβ (42/40) ratio [t(7)=2.2, P=.03], but had no effect on Tau expression (**Figure 6B**). Elevated Aβ42/40 ratio in the PFC is indicative of *Alzheimer’s pathologic change*(e.g., A+T-(N)-) as reflected by amyloid pathology (**Table 2**), which is thought to occur at a (moderate) stage of AD (Braak and Braak, 1998); thus, alcohol drinking appears to enhance the onset of PFC-linked AD pathology in 3xTg-AD mice.

#### Medial Prefrontal Cortex (mPFC) and Amygdala (AMY)

Alcohol had no effect on the Aβ (42/40) ratio in the mPFC or AMY. However alcohol drinking was associated with an increase in total Tau expression in both brain regions: mPFC - [t(7)=3.0,P=0.02] (**Figure 6C**); AMY - [t(7)=3.6, P=0.009] (**Figure 6D**). This result is consistent with *Non-AD pathologic change* (e.g., A-T-(N)+, **Table 2**); however, Tau-related pathology is associated with late-stage AD suggesting that alcohol may exacerbate the onset of pathology in the AMY.

#### Additional brain regions

No changes in Aβ42/40 ratio or total Tau protein were detected in samples from nucleus accumbens (Acb), medial hippocampus (MHPC), lateral hippocampus (LHPC), CA1 subregion of the hippocampus (CA1), and the medial entorhinal cortex (MEC; data not shown.

#### Conclusions

<u>First</u>, based on evidence that 6 – 8 month old 3xTg-AD mice exhibit AD-like pathology expressed as increased Aβ in the HPC, cortex, and AMY (Oddo et al., 2003b), these data suggest that alcohol increased the magnitude of AD-like pathology in the LEC and PFC during the initial cortical stages (**e.g, Figure 2**). <u>Second</u>, 3xTg-AD mice are known to exhibit Tau pathology at 12-15 months; thus, the appearance of elevated total Tau in LEC, mPFC and AMY at 8-months of age suggests that alcohol drinking promoted rapid onset of pathology in these brain regions. <u>Third</u>, it is important to note that these data were obtained 1-month after alcohol use suggesting the presence of a long-term negative impact on AD-like pathology. Moreover, by focusing on biomarkers within the NIA-AA Research Framework, these findings integrate with the broader field of age-related neurodegeneration and suggest that alcohol use is a risk factor for AD-like pathology in vulnerable individuals. Increased knowledge of specific pathologies associated with alcohol use in AD has the potential to aid diagnosis and treatment.

### Tau pathology: Hyperphosphorylation of Tau at GSK-3β site in 3xTg-AD mice (1-month post alcohol)

The presence of paired helical fragments formed by hyperphosphorylation of microtubule-associated protein Tau is a hallmark pathology of AD (Raskin et al., 2015). The Tau protein can be phosphorylated at a combined total of 85 serine, threonine, and tyrosine residues that constitute about 20% of the total protein (Tenreiro et al., 2014); however, there appears to be specificity with respect to disease progression. In late stage AD (Braak stage V/VI; Severe dementia, anxiety, depression) phosphorylation of a cluster of Tau residues termed AT8 (Ser199/Ser202/Thr205) is elevated in cortical brain tissue from human AD patients from 4- to 13-fold over controls (Neddens et al., 2018). Strikingly, analysis of the transentorhinal cortex showed that Tau phosphorylation at Ser199 was elevated as much as 160-fold over healthy controls suggesting a high degree of relevance for the Ser199 residue in disease progression (Neddens et al., 2018).

Although Tau is phosphorylated by numerous kinases, evidence indicates that glycogen synthase kinase-3 beta (GSK-3β) phosphorylation at Ser199 and Ser202 (pTau-Ser199/202) is a key component of AD pathology and thought to be a viable therapeutic target (Llorens-Martin et al., 2014). Interestingly, GSK-3β is an alcohol-sensitive gene (Wolen et al., 2012) and phospho-protein (Cheng et al., 2017; Liu et al., 2017) in a variety of brain regions, including the HPC of adult mice exposed to prenatal alcohol (Cunningham et al., 2017). Therefore, we asked if Tau phosphorylation at the GSK-3β (Ser199/202) site is altered by alcohol drinking in 3xTg-AD mice, which would indicate a potential target of alcohol.

3xTg-AD and WT mice were sacrificed 1-month after alcohol or saccharin drinking (as described above) for qualitative assessment of pTau-Ser199/202 (pTau) immunoreactivity (IR). pTau -Ser199/202 IR was not observed in WT mice (**Figure 7A**). By contrast, alcohol produced pronounced hyperphosphorylation of Tau-Ser199/202 IR in dorsal hippocampal neuronal cell bodies and projections of 3xTg-AD mice as compared to saccharin controls (**Figure 7A**). No difference was observed in BLA of 3xTg-AD mice (**Figure 7B**).

**Figure 7.**
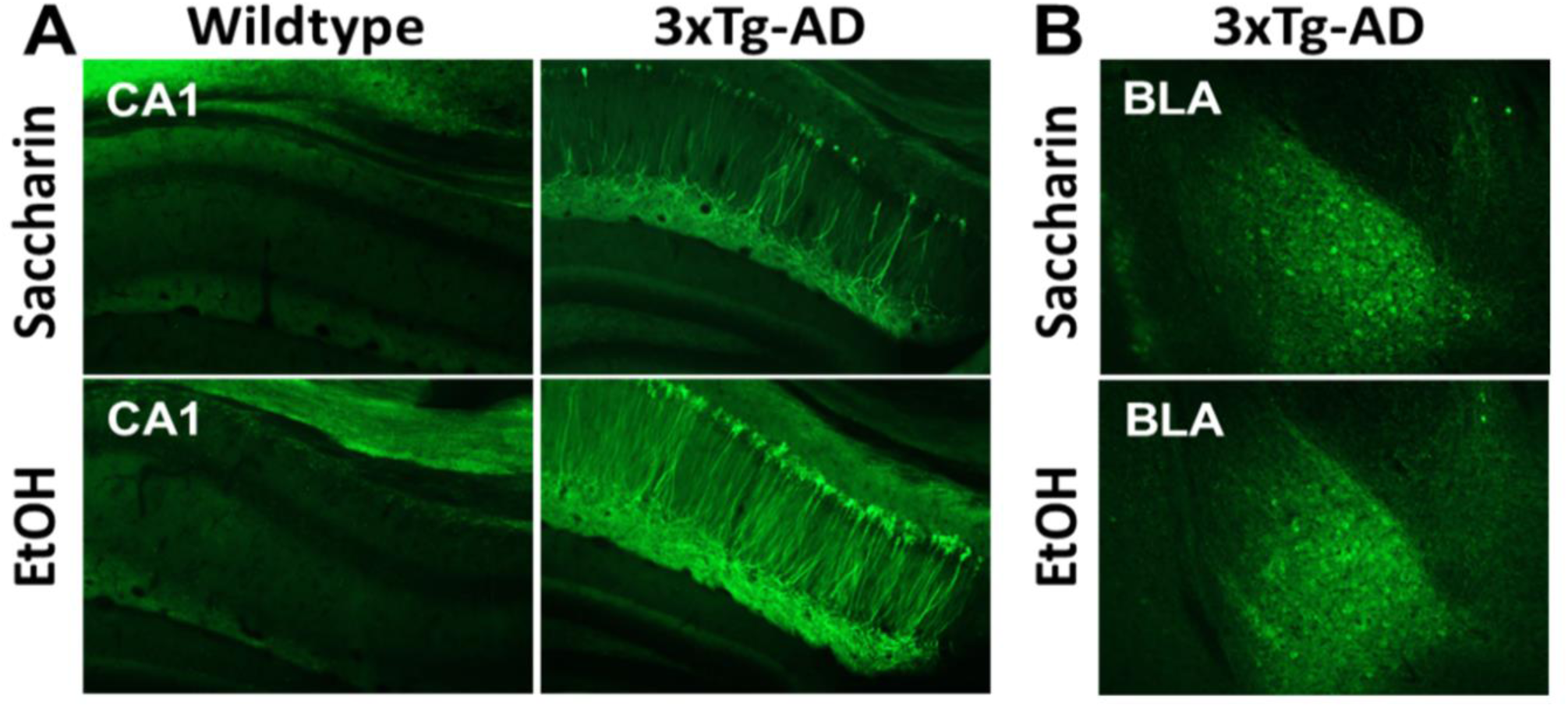
pTau-Ser199/202 immunoreactivity (IR) in WT and 3xTg-AD mice 1-month post alcohol or saccharin drinking. (**A**) Coronal sections of dorsal hippocampus showing prominent IR in neuronal cell bodies and projections of CA1 only in 3xTG-AD mice with upregulation post alcohol (EtOH). (**B**) Basolateral amygdala (BLA) sections showing no alcohol-induced difference in pTau-Ser199/202 IR. Images acquired on Olympus BX51 microscope at 10x magnification.

The results of this study are consistent with prior findings in 3xTg-AD mice showing age-dependent increases in Tau AT8 expression. Untreated 3xTg-AD mice show low levels of pTau AT8 immunoreactivity the HPC (CA1 region) at 6-months of age but prominent expression in neurons and projections at 15-months of age (Oddo et al., 2003a). This age-dependent progression of AT8 expression is similar to the present result showing that alcohol drinking (1-month post exposure) was associated with hyperphosphorylation of Tau at the Ser199/202 residues in the HPC. As noted above, Tau phosphorylation at these sites in humans is associated with advanced Braak stage V/VI Alzheimer’s disease that is characterized by severe dementia and other neuropsychiatric symptoms. This raises the possibility that alcohol-induced upregulation of GSK-3β phosphorylation of Tau (Ser199/202) may mediate hippocampal-dependent behavioral pathology associated with alcohol use. Indeed, the present results show that alcohol exposed 3xTg-AD mice have deficits in spatial memory (**Figure 5A**) and sensorimotor gating (PPI, **Figure 5B**), which are both modulated by hippocampal activity. It is also interesting to note, that there was no difference in pTau (Ser199/202) IR in the BLA suggesting that apparent emotional dysregulation shown here in fear conditioning (**Figure 5C**) is not related to pTau (Ser199/202) pathology. Elucidating the impact of alcohol use on Tau phosphorylation in the hippocampus has potential to move the field forward in understanding behavioral pathologies that occur in older individuals and may recommend therapeutic strategies for alleviation of specific cognitive deficits in patients with a history of alcohol use.

### Analysis of Akt/mTOR phosphoprotein pathway throughout the brain after chronic alcohol: Decreased expression of multiple mTOR/Akt phosphoproteins in 3xTg-AD alcohol exposed mice (1-month post alcohol)

To gain further understanding of the long-lasting impact of chronic non-dependent alcohol drinking on molecular mechanisms of AD, we assessed the phosphorylation state of the Akt/mTOR cell signaling pathway. AKT/mTOR signaling regulates a wide array of molecular and cellular functions in the brain including nutrient uptake, glucose metabolism, cell growth and survival, transcription and migration (Yu and Cui, 2016). Importantly, the Akt/mTOR pathway is implicated in AD pathology in humans and animal models (Oddo, 2012) and alcohol addiction (Neasta et al., 2014). For example, glycogen synthase kinase-3 (GSK3) is critical to the hyper-phosphorylation of Tau, has been linked to increased Aβ production, and Aβ mediated cell death (Avila et al., 2012; de la Monte et al., 1999; Hernandez, 2013; Hooper et al., 2008). GSK3 also regulates the positive reinforcing effects of alcohol and is sensitive to acute alcohol treatment and withdrawal (Faccidomo S. et al., in press; Neasta et al., 2011; van der Vaart et al., 2018b). Additional proteins in the mTOR/Akt pathway such as Akt, PTEN, p70S6K, RPS6, ERK_1/2_ and mTOR are targets of interest in understanding the pathogenesis of AD as this cell signaling pathway is an important modulator of cell growth and macroautophagy maintaining a balance critical to the health of cells (Jung et al., 2010; Norambuena et al., 2017; Pei and Hugon, 2008a). Overall, altered function of the Akt/mTOR pathway has been documented following acute and chronic alcohol exposure (Besheer et al., 2012; Faccidomo et al., 2015b; Fu et al., 2016; Hagner et al., 2009; Neasta et al., 2010; Spanos et al., 2012) and implicated in neural and behavioral pathology associated with neurodegeneration, suggesting that this cell signaling pathway may be a target of alcohol in the development and expression of AD.

In the present study, the brains of 3xTg-AD mice were probed for changes to phosphoproteins of the mTOR/Akt pathway after chronic, non-dependent alcohol [(25% w/v)+saccharin (0.1% w/v) vs. water (n=5)] as compared to saccharin [(0.1% w/v) vs. water (n=5)] drinkers in to our standard 2-bottle choice 24-h home-cage method. Brains were removed approximately 1 month after the final alcohol or saccharin drinking session (at 8 months of age), so the following results indicate lasting changes provoked by the interaction of AD and chronic alcohol drinking. Both groups of mice drank similar volumes and alcohol drinking mice consumed an average of 15.7 g/kg/day representing a moderate level of alcohol consumption, but not reaching binge or heavy drinking levels. Overall, the results indicate dysfunction of the mTOR/Akt phosphoprotein signaling with the predominate trend showing decreases throughout the brains of 3xTg-AD mice with a history of alcohol drinking.

#### Hippocampus

In 3 subregions of the HPC, (medial, lateral, and CA1) analyzed in this study, the effects on the mTOR/Akt pathway were sub-region specific. This is an important distinction from other researchers looking at hippocampal changes as the subregion is not often specified. In the **MHPC**, no significant changes were observed in the expression of Akt/mTOR phosphoproteins between alcohol and saccharin exposed 3xTg-AD mice (**Table 3**). In the **LHPC**, a history of alcohol drinking in 3xTg-AD mice was associated with significant reductions in phosphorylation of IRS1 t(6)=1.974, P=0.048)] and P70S6K [t(6)=2.078, P=0.042)] as compared to saccharin controls (**Table 3**). Finally, in **CA1**, a history of chronic alcohol drinking in 3xTg-AD mice was associated with a significant reduction in phosphorylated mTOR [t(7)=3.011, P=0.01)] and PTEN [t(7)=3.021, P=0.01)] as compared to saccharin controls (**Table 3**).

**Table 3.** mTOR/Akt Phosphoprotein cell signaling in 3xTg-AD mice. This table shows the percent change in relative protein expression of 3xTg-AD alcohol drinking mice compared to saccharin drinking mice at 1-month post alcohol (8-months old). Relative change calculated using β-Tubulin as loading control. All analytes and brain regions were treated as independent samples for analysis. Shaded boxes with * indicate P<0.05; for one-tailed, t-test. Abbreviations: protein kinase B (Akt/PKB), mitogen-activated protein kinase/extracellular signal-regulated kinases pathway (ERK 1/2), glycogen synthase kinase 3α (GSK3α), glycogen synthase kinase 3β (GSK3β), insulin-like growth factor 1 receptor (IGF1R), insulin receptor (IR), insulin receptor substrate 1 (IRS1), mammalian target of rapamycin (mTOR), ribosomal protein s6 kinase β-1 (p70S6K), phosphatase and tensin homolog (PTEN), ribosomal protein S6 (RPS6), mTOR complex 2 (TSC2)

| Protein | Phosphorylation site | mPFC | PFC | Acb | AMY | MHPC | LHPC | CA1 | LEC | MEC |
| --- | --- | --- | --- | --- | --- | --- | --- | --- | --- | --- |
| **Akt/PKB** | (Ser473) | 111.3 | 91.4 | 90.7 | 95.7 | 113.5 | 103.2 | 81.7 | 68.6 | 81.8 |
| **ERK 1/2** | (Thr185/Tyr187) | 101.9 | 80.3 | 91.4 | **69.0\*** | 109.6 | 74.3 | 93.0 | 74.0 | 64.7 |
| **GSK3α** | (Ser21) | 72.6 | 71.1 | 91.4 | 97.5 | 82.2 | 84.9 | 122.8 | 52.4 | **60.5\*** |
| **GSK3β** | (Ser9) | 130.6 | 99.4 | 90.6 | 96.8 | 114.9 | 91.7 | 83.4 | 101.2 | 109.6 |
| **IGF1R** | (Tyr1135/1136) | 89.7 | 86.2 | 91.5 | 86.7 | 79.7 | 68.2 | 104.7 | **59.7\*** | **69.4\*** |
| **IR** | (Tyr1162/1163) | 105.6 | 113.2 | 81.9 | 86.8 | 98.8 | 91.1 | 92.5 | **54.4\*** | 80.4 |
| **IRS1** | (Ser636) | 94.0 | 88.3 | 82.7 | 106.4 | 80.0 | **66.3\*** | 88.2 | 56.4 | **64.3\*** |
| **mTOR** | (Ser2448) | 102.0 | 86.6 | 82.8 | 74.0 | 84.1 | 85.9 | **76.3\*** | 68.5 | 82.2 |
| **p70S6K** | (Thr389/412) | 74.7 | 117.1 | 50.7 | 87.7 | 65.1 | **49.4\*** | 142.0 | 62.0 | 59.7 |
| **PTEN** | (Ser380) | 104.6 | 106.1 | 89.1 | 94.7 | 108.3 | 113.6 | **80.3\*** | **72.9\*** | 109.1 |
| **RPS6** | (Ser235/236) | 96.1 | 93.1 | 81.3 | 93.9 | 87.7 | 83.7 | 103.5 | 60.2 | **73.7\*** |
| **TSC2** | (Ser939) | 104.8 | 117.0 | 99.7 | 119.3 | 115.4 | 78.6 | 81.3 | 94.1 | 101.4 |

These results indicate that a history of alcohol drinking is associated with downregulation of Akt/mTOR phospho-signaling in sub-regions of the HPC in 3xTg-AD mice. The impact of alcohol was selective to the LHPC (IRS1 and p70S6K) and CA1 (mTOR and PTEN). This is consistent with evidence showing that levels of p-mTOR (Ser2448) and p70S6K (Thr389) are decreased in the cortex of double APP/PS1 transgenic mice as compared to WT controls (Lafay-Chebassier et al., 2005). Similarly, treatment of SH-SY5Y with 20 μM Aβ decreases p-mTOR (Ser2448) expression, and pharmacological inhibition of mTOR induced cell death (Lafay-Chebassier et al., 2006). However, other reports show increased mTOR or PTEN in AD and mixed results in the alcohol literature (Caccamo et al., 2018; Fu et al., 2016; Griffin et al., 2005; Hagner et al., 2009; Li and Ren, 2007; Norambuena et al., 2017; Oddo, 2012; Tramutola et al., 2015; Wang et al., 2014). Thus, it will be critical for future studies to carefully parse out changes in these phosphoproteins due to AD pathology, long-term alcohol, and normal aging.

In the 3xTgAD mice, prior studies have found decreased levels of IRS-1 in the membrane fraction of hippocampal lysates (Ma et al., 2009). Moreover, phosphorylated IRS-1 is increased in 9-month old 3xTg-AD mouse HPC and cortex but decreased at 10- and 15-months of age (Velazquez et al., 2017) when neural and behavioral pathology is most severe. Thus, the reduction in hippocampal IRS-1 phosphorylation seen in 8-month old 3xTg-AD mice in the present study suggests that alcohol drinking may have exacerbated the onset or magnitude of AD-like pathology. This effect of alcohol was measured 1-month after exposure but is consistent with reduced IRS-1 phosphorylation seen after ethanol (100 mM) in cell culture (Xu et al., 1995).

In previous studies of AD, P70S6K is more often reported to be increased (An et al., 2003; Jin-Jing, 2008); however, alcohol has been reported to decrease P70S6K like it has been observed here (Hagner et al., 2009; Li and Ren, 2007), but to increase with acute exposure (Fu et al., 2016). Because this comparison is limited to saccharin or alcohol drinking 3xTg-AD mice, future studies will have to reconcile the differential effects on the phosphorylation of the P70S6K protein and the magnitude of the effects compared to wildtype mice.

#### Entorhinal cortex

The entorhinal cortex is sensitive to early AD pathology and this is reflected in both the brain and behavior. It is relevant in AD pathology in part because of a major pathway projecting from the AMY to the entorhinal (primarily lateral) as well as a substantial pathway projecting from the entorhinal cortex to the HPC. However, to date, the only study that has considered the role of the mTOR pathway in the AD pathology of the entorhinal cortex did so by pharmacologically inhibiting mTOR via chronic systemic treatment with rapamycin (Siman et al., 2015). This group observed a decrease in Tau-induced neuronal loss in the LEC, but not total Tau or mRNA levels. Our results suggest notable dysfunction of both the LEC and MED mTOR/Akt phosphoproteins after chronic alcohol consumption. In the **LEC**, alcohol drinking significantly decreased IGF1R [t(8)=1.905,P=0.047)], IR [t(8)=2.211,P=0.029)], and PTEN [t(8)=1.976,P=0.042)]; while in the **MEC**, GSK3α [t(7)=2.590,P=0.018)], IGF1R [t(7)=2.949,P=0.011)], IRS1 [t(7)=3.249,P=0.007), and RPS6 [t(7)=2.104,P=0.037)] were all significantly reduced when compared to saccharin drinkers.

#### Amygdala

In the **AMY**, a history of alcohol drinking was associated with a significant decrease in pERK1/2/MAPT1/2 [t(7)=1.905, P=0.049)]. No other significant changes were observed; however, trends toward a reduction were observed subregions of the HPC and entorhinal cortex (**Table 3**).

This finding is consistent with prior work showing that ERK1/2 phosphorylation is reduced in the AMY following alcohol withdrawal and associated with increased anxiety-like behavior in rats (Pandey et al., 2008). In the present study, 3xTg-AD showed a similar pattern of results with reduced pERK1/2 expression in the AMY associated with enhanced cued fear response. Thus, dysregulation of ERK1/2 activity in the AMY may mediate heightened anxiety and fear related to AD pathology. Other evidence, however, shows that pERK1/2 is increased in the AMY of APP_Ind_ transgenic mice immediately after cued fear conditioning, suggesting that ERK signaling regulates enhanced fear response associated with AD (Espana et al., 2010). Interestingly, the observed decrease in pERK1/2 observed in the AMY of alcohol exposed 3xTg-AD mice is similar to the effects of chronic unpredictable stress (Chandran et al., 2013). Thus, it will be important for future studies to evaluate pERK1/2 following cued fear conditioning in alcohol exposed 3xTg-AD mice to determine if the observed basal downregulation is associated with insensitivity, or compensatory increased sensitivity, of ERK signaling under stressful conditions.

#### Conclusions

The mTOR/Akt pathway is critical in the regulation of cell growth, proliferation, and macroautophagy. The pathway is inextricably linked to aging and is regulated by growth factors, glucose, insulin, and stress. Aberrant signaling of the mTOR/Akt pathway has been implicated in human studies of AD and in numerous animal models, though the literature isn’t consistent about specific increases or decreases of total or phosphorylated proteins. To account for this discrepancy, it has been proposed that Akt/mTOR phosphorylation is decreased in neurons that are sensitive to AD and increased or maintained in neurons that are insensitive to AD (Pei and Hugon, 2008b). Consistent with this premise, the present study found evidence for no change, significant decreases, and trends for increased phosphorylation in the Akt/mTOR pathway that was protein- and brain region-dependent. Moreover, the present study evaluated phosphorylation of Akt/mTOR signaling proteins in relatively young 3xTg-AD mice (8 months of age) at 1-month after alcohol exposure, which may reflect direct long-lasting effects of alcohol or compensatory changes. For these reasons, we suggest that understanding the impact of alcohol drinking on the Akt/mTOR signaling pathway in AD will require delineation of age-, time-, and brain region-dependent effects of alcohol. It is plausible that vulnerability to AD-like neural and behavioral pathology will vary as a function of one or more of these variables.

Likewise, the Akt/mTOR pathway is both directly and indirectly impacted by alcohol consumption via its direct inhibition of IRS1, and indirectly through modulation of metabolic function. It is for these reasons invariably complicated to study the interaction of AD and alcohol consumption on the mTOR/Akt pathway, but also imperative. The ability of multiple proteins in the mTOR/Akt pathway to directly affect the production of AΒ and hyperphosphorylation of Tau makes it a potential therapeutic target for early disease.

## GENERAL DISCUSSION

Alcohol use impacts health and disease across the lifespan. Especially relevant to the aging population, emerging evidence suggests that alcohol use is a potential risk factor for AD and other forms of dementia. The purpose of the present study was to test aspects of the overall hypothesis that alcohol use exacerbates the onset and magnitude of AD-like pathology.

The scientific premise of this work was based primarily on our prior analysis of the neuroproteome that revealed a striking, and unanticipated, major association between alcohol and AD pathology in C57BL/6J mice (Agoglia and Hodge, 2017; Salling et al., 2016). A newly conducted bioinformatics analysis of those data identified three primary mechanisms of AD (MAPT, PSEN-1, and APP proteins) as the statistically most likely upstream modulators of alcohol-sensitive protein networks in the PFC and amygdala. Based on this predicted relationship, we sought to evaluate the impact of voluntary nondependent alcohol drinking on AD-like neural and behavioral pathologies in the 3xTg-AD mouse model of AD, which expresses human MAPT, PSEN-1, and APP transgenes (Oddo et al., 2003b).

### Alcohol-induced Behavioral Deficits

As noted above, AD is characterized by a progressive decline in cognition (e.g., memory, stimulus processing) and other behavioral functions (e.g., anxiety, fear). Importantly, cognitive decline in conjunction with comorbid neuropsychiatric conditions such as anxiety may reflect early stages of AD pathology (Buckley et al., 2016; Jessen et al., 2014). Evidence indicates that 3xTg-AD mice exhibit a variety of behavioral phenotypes consistent with progressive cognitive decline and altered emotional state in a manner that is associated with development of underlying neural pathology (Filali et al., 2012; Huber et al., 2018; Pietropaolo et al., 2014; Romano et al., 2014; Webster et al., 2014).

The behavioral test battery used in the present study showed that a history of alcohol drinking (4-months) was associated with deficits in 3xTg-AD mice as compared to saccharin exposed and WT controls. As discussed above, 3xTg-AD mice, and other mouse models of AD, develop age-dependent pathologies in these behavioral domains irrespective of alcohol exposure (Webster et al., 2014). However, at the age range tested (7 – 8 months) saccharin exposed 3xTg-AD mice in the present study did not show major behavioral abnormalities as compared to wildtype controls with the only notable deficit appearing in spatial learning in Morris Water Maze (e.g., acquisition). By contrast, a history of nondependent alcohol drinking produced behavioral deficits in spatial memory as tested in the Morris Water Maze (probe trial), sensorimotor gating (PPI of the startle response), and emotional processing as indexed by enhanced cued fear response.

Both male and female mice were tested in the present study, but sample size did not allow statistical evaluation of sex as a biological variable. That said, we observed tendencies for sex-based differences in Morris Water Maze and prepulse inhibition performance with female data trending toward a greater deficit. This raises the possibility that females may show enhanced vulnerability to the age-dependent effects of alcohol on cognitive function. However, results also showed a trend for female mice to consume higher doses of alcohol than males, which is consistent with evidence from a wide sample of rodent studies (Becker and Koob, 2016); thus, it will be necessary in future preclinical studies to equate alcohol intake between sexes using between group yoking procedures. In addition, sex-based differences in alcohol pharmacokinetics may account for differential neural or behavioral effects, which complicates direct dose-dependent comparisons. Nevertheless, it is important to note that the impact of alcohol on cognitive (memory) function in older adults is more pronounced in women (Lewis et al., 2019) underscoring the importance of conducting future preclinical work to elucidate neural mechanisms that might contribute to female vulnerability.

Overall, these data show that voluntary alcohol drinking in 3xTg-AD mice is associated with impaired cognitive and emotional functions as compared to saccharin and wildtype controls, which mimic behavioral pathologies seen in humans with AD and suggest that alcohol exacerbated the onset of these symptoms. However, there remains a gap in knowledge regarding the sex- and age-dependent impact of alcohol use on AD-like pathology in older individuals. To address this question, future preclinical studies could evaluate the impact of alcohol exposure in 3xTg-AD mice at timepoints that correspond pre- and post-pathology phases of development in male and female mice.

### Impact of Alcohol on AD Biomarkers in Specific Brain Regions

As discussed above, the jointly proposed NIA-AA Research Framework calls for a biological definition of AD with emphasis on categorizing AD pathology based on measurement of specific biomarkers including Aβ, Tau, neurodegeneration, and cognitive symptoms (Jack et al., 2018). In this framework, specific combinations of individual measures have different interpretations. Presence of Aβ biomarkers (e.g., the Aβ 42/40 ratio) determines presence on the Alzheimer’s continuum (Table 2). Pathologic tau biomarkers (e.g., pTau) can indicate non-AD pathology if detected in isolation; however, combined Aβ and Tau biomarkers indicates presence of AD because both are required for diagnosis of the disease. The addition of cognitive symptoms is also used for disease staging with dementia (e.g., cognitive impairment in several domains with additional behavioral symptoms) indicating the most severe stage (Jack et al., 2018). In this study, we took a translational approach and evaluated AD biomarkers within specific brain regions of the 3xTg-AD mice and evaluated the results in the context of the NIA-AA framework.

Results from the present preclinical study indicate that a history of alcohol exposure is associated with a biomarker profile that is consistent with alcohol-induced expression of Alzheimer’s disease in 3xTg-AD mice. First, alcohol exposure resulted in an increase in the Aβ (42/40) ratio in the LEC and PFC (**Figure 6A and 5B**). Aβ (42) is the toxic form of Aβ and is upregulated in postmortem brains of AD patients. This results in an A+ categorization and is indicative of presence on the AD continuum (Table 2). Second, we observed an increase in total Tau protein expression in the LEC, mPFC and AMY. An increase in total Tau protein is indicative of neurodegeneration or neuronal injury and represents an N+ categorization. These data indicate that alcohol use was associated with an A+T(N)+ combination, which indicates Alzheimer’s and concomitant non-Alzheimer’s pathology in the LEC, which is an initial anatomical target of AD pathology (Table 2). In addition, immunohistochemistry was used to measure pTau expression. Qualitative results showed a prominent upregulation of pTau (Ser199/202) in the hippocampus (CA1) of alcohol exposed 3xTg-AD mice. Taken together across brain regions, these data suggest that alcohol exposure produced an A+T+(N)+ biomarker profile, which is consistent with Alzheimer’s disease.

It is a limitation of the present study that we did not measure pTau in the multiplex immunoassay. This was due to a limitation of the technique, which could not differentiate and count individual microbeads labeled with pTau due to excessive protein aggregation and bead clustering. Methodological development efforts are underway in our laboratory with the goal of disaggregating pTau prior to measurement; however, future studies will utilize immunoblot or immunohistochemistry approaches to obtain pTau measurements across the brain.

Overall, these results suggest that alcohol use is a modifiable risk factor for AD. By focusing on AD biomarkers within the NIA-AA Research Framework, this work integrates with the broader field of age-related neurodegeneration and extends efforts to elucidate the impact of alcohol age-related neurodegeneration. Increased knowledge of specific pathologies associated with alcohol use in AD has potential to aid diagnosis and treatment.

### Alcohol-induced Hyperphosphorylation of Tau in the Hippocampus

As noted above, glycogen synthase kinase 3 (GSK-3) is a serine-threonine kinase that is expressed in the mammalian brain system in two isoforms, GSK-3α and GSK-3β (Woodgett, 1990). Dysregulated GSK-3β activity is associated with several neurological and neuropsychiatric diseases including Alzheimer’s (Cai et al., 2012) and is thought to serve as a molecular link between Aβ and Tau pathology (Llorens-Martin et al., 2014). GSK-3β cleaves APP to Aβ and directly phosphorylates Tau at multiple amino acid residues including Ser199/Ser202. Acute and chronic alcohol exposure is known to increase GSK-3β phosphorylation (Neasta et al., 2014; Neasta et al., 2010; Neasta et al., 2011; van der Vaart et al., 2018a) but the long-term effects of alcohol use on GSK-3 activity and function remain to be elucidated. Results from this study show that a history of alcohol use (1-month post alcohol) is associated with hyperphosphorylation at the GSK-3 site on Tau (Ser199/Ser202) in hippocampus (CA1) subregion of 3xTg-AD mouse brain.

The heightened pTau (Ser199/202) immunoreactivity observed in 3xTg-AD mice suggests that nondependent alcohol use may produce enduring changes in the functional effects of GSK-3 that underly neural pathology. Moreover, GSK-3 may represent a therapeutic target for neurodegeneration associated with alcohol use or abuse. A growing literature indicates that pharmacological or genetic inhibition of GSK-3 blocks cleavage of APP, which reduces Aβ deposits and restores hippocampal-dependent spatial memory (water maze) in mice (King et al., 2014; Ly et al., 2013; Rockenstein et al., 2007). Similarly, chronic GSK-3β inhibition reduces Tau phosphorylation and improves memory in 3xTg-AD mice (Huang et al., 2019). Although GSK-3 inhibitors have produced mixed results in clinical trials, selective non-ATP competitive compounds are well-tolerated in vivo and serve as useful experimental tools to probe mechanism of action and may have therapeutic efficacy under specific pathological conditions. For example, non-ATP competitive TDZD derivative tideglusib (200 mg/kg, p.o. for 3 months) reduces AD-like pathology in mice including decreased Tau phosphorylation and Aβ deposition in the entorhinal cortex and HPC (CA1) with associated prevention of memory deficits in the Morris Water Maze (Sereno et al., 2009). More research is needed to determine if GSK-3 inhibition modulates Tau pathology seen following alcohol exposure.

### Impact of Alcohol Use on the Akt/mTOR Pathway

Results of this study showed a general downregulation of multiple Akt/mTOR phosphoproteins in AMY, LHPC, CA1, LEC, and MEC suggesting that a history of alcohol use produced an enduring general disruption of Akt/mTOR signaling in brain regions associated with AD pathology. As noted above, the Akt/mTOR signaling pathway regulates cell growth and proliferation, as well as macroautophagy. Given the neurodegenerative nature of AD and chronic alcohol consumption, it is not unexpected that together they would disrupt these basic processes and, potentially, accelerate decline. However, the mechanisms that are specifically responsible are harder to understand as the nature of phosphorylated proteins is only one change in response to environmental demands on both a micro and macro scale in any signaling pathway.

GSK-3 is a key member of the Akt/mTOR cell signaling pathway. Given the upregulation of pTau (Ser199/202) immunoreactivity observed in CA1, we predicted that alcohol use would be associated with altered GSK-3 phosphorylation in this brain region. However, alcohol exposure was associated with trends for increased GSK-3 α (Ser21) and decreased GSK-3β (Ser9) phosphorylation in CA1. Since GSK-3α activity is increased by phosphorylation and GSK-3β activity is increased by dephosphorylation, these trends in phosphorylation may have been enough to support increased Tau phosphorylation. Further work is needed to determine if alcohol-induced changes in GSK-3 phosphorylation modulate Tau phosphorylation and expression of AD-like pathology.

Understanding the delicate balance of the various proteins in the Akt/mTOR pathway in relation to alcohol’s impact on AD could ultimately provide novel targets for therapeutic strategies that are able to restore balance rather than simply increase or decrease specific proteins. It should also be acknowledged that there is a need for future studies to consider the impact of changes in the periphery (ie, liver, pancreas) on the CNS. Changes in the brain, especially to regulatory processes, result in changes in the periphery that can in turn act as a feedback loop to result in further changes to the regulatory processes in the brain. Alcohol is well known to have detrimental impact on various organs and these changes in metabolism and insulin regulation are more than likely also factors that need to be considered when studying the CNS mTOR/Akt pathway in the context of AD and long-term alcohol consumption.

### Alcohol-induced Impact on the Neuroproteome and AD

It is noteworthy that many of the alcohol-sensitive proteins that were significantly changed in our proteomic networks are also associated with major neurodegenerative diseases such as AD and schizophrenia (Hensley et al., 2011; Martins-De-Souza et al., 2010). The DPYSL protein, which was significantly downregulated in the amygdala network, stands out for many reasons and may be an ideal therapeutic target for AD. The gene DPYSL encodes a protein called collapsin response mediator protein 2 (CRMP2) which is a member of a family of CRMP proteins that regulate microtubule formation and axonal growth (Charrier et al., 2003). CRMP is highly expressed in the CNS and it is important for microtubule formation and stability, vesicle trafficking and ion channel stability, among other functions. There has been exponential interest in the functional role of this protein in disease pathology because it is hyperphosphorylated in the brains of both AD patients and it is known to be a key phosphorylation substrate for GSK-3β, CDK5 and CAMKII (Soutar et al., 2009). Phosphorylation of CRMP at these sites renders it unable to function properly and is proximally associated with plaque and tangle formations in AD brains (Yoshida et al., 1998). Further, it has been shown that CRMP2 co-aggregates with Tau in 3xTg mice at an early age, suggesting that CRMP could be used as an early biomarker for AD, prior to expression of significant behavioral deficits (Cole et al., 2007).

Our behavioral data suggest that 3xTg mice show impaired spatial learning (MWM acquisition) by 7 – 8 months of age and that a history of moderate alcohol drinking leads to impaired short-term spatial memory (probe test). Likewise, CRMP1 KO mice show decreased LTP in CA1 and impaired acquisition of spatial learning in the Morris Water Maze as well as impaired short-term memory on a probe (Su et al., 2007). Together, these data suggest that the CRMP family of proteins may regulate some of the behavioral deficits that are observed in 3xTg mice will high likelihood of contributing to the molecular deficits. Because of its potential usefulness as an early biomarker for AD, CRMP may be a logical therapeutic target for reversing or mitigating some of the pathological and behavioral deficits associated with AD. Indeed, this hypothesis was tested by Hensley and colleagues in 3xTg mice (Hensley et al., 2013). They administered a drug that binds to CRMP2 and found that it diminished the consequences of disease progression as measured by decreased latency to find a platform in the MWM and a decrease in AB and APP immunoreactivity in the HPC. Alternatively, pharmacologically increasing expression of CRMP2 has been proposed as a possible therapeutic because it could counteract the neuronal stress associated with AD. The atypical anti-depressant, tianeptine, has a multitude of pharmacological actions, and mechanistically, it has been shown to interfere with CAMKII-GluA1 binding and pharmacologically increase CRMP expression (Hensley et al., 2011). This drug is widely prescribed for depression, yet its mechanisms of action suggest that it could be an effective candidate for the treatment of AD. Importantly, we have recently published evidence that CRMP is more abundant in the adult vs. adolescent PFC proteome (Agoglia et al., 2017), we show here that CRMP is downregulated by alcohol drinking and pharmacological treatment with tianeptine can indeed reduce levels of binge drinking in adult mice (Agoglia et al., 2015b). Future studies might investigate whether tianeptine could be effective in reversing and/or ameliorating the AD behavioral and brain pathology induced by chronic alcohol drinking.

A striking commonality of the alcohol-sensitive proteins that were upregulated in the mPFC and AMY is their notable relevance to AD pathology. Creatine kinase B (CKB), hexokinase (HK1), syntaxin binding protein 1 (STXB1/MUNC-18), amphiphysin (AMPH), complexin, and heat shock proteins are all upregulated in the cortex of AD patients and in normal, alcohol-drinking C57BL/6J mice (Donovan et al., 2012; Jacobs et al., 2006; Lynn et al., 2010; Rivera et al., 2018). Further, CKB and HK1 are more prominently increased in the brains of individuals with early- vs late-stage disease suggesting that proteins relevant for energy use and metabolism may be early markers of brain pathology and stress. Munc-18 and complexin are part of a family of neuronal vesicular proteins regulating the SNARE complex which is involved in rapid exocytosis and neurotransmitter release at neuronal synapses (Pabst et al., 2002; Sudhof, 2013; Tang et al., 2006). Mechanistically, this complex directly binds to APP causing increased aggregation of Aβ in the brain and indeed, these proteins are physically located in proximity to aggregated Aβ (Chaufty et al., 2012; Jensen et al., 2018).

Thus, although alcohol drinking induced differential changes in the PFC and AMY proteome, specific target proteins in each brain region are known to regulate aspects AD pathology. Moreover, identified networks or protein groupings were identified by IPA as likely to be downstream of APP, PSEN-1 and Tau, which suggests collective involvement of unique protein networks within the PFC and AMY in AD. This indicates that alcohol use may influence AD pathology via distinct but functionally related protein networks that vary across brain regions. Therefore, efforts to more fully elucidate the impact of alcohol use on AD will benefit from a systems approach that delineates adaptations in specific molecular mechanisms within brain regions, but also considers common functional effects of distinct proteins between brain regions.

### Conclusions and Future Directions

Alcoholism and Alzheimer’s Disease are both neuroinflammatory and neurodegenerative disorders and as such, may share an overlapping etiology. The data show that indeed, the neuropathological changes that are associated with both disorders can be linked to mechanisms related to AB, Tau and the mTor pathway. The relevance of our alcohol proteomic networks to AD indicate that there may be important systems that could be useful early biomarkers (CRMP, HSK, AMPH, GSK) for identifying at-risk populations that could be then targeted with medications (tianeptine, lacosamide, tideglisib) with prophylactic treatment to prevent or minimize the development of AD. Given the strength of these associations, further research in both clinical and pre-clinical populations to test these hypotheses is warranted. Together, these data strongly suggest that some (or all) of these molecular mechanisms may be dysregulated by moderate alcohol use and that this may increase risk for developing and or exacerbating AD in susceptible populations.

Preclinical strategies, such as those reported here, allow collection of new and important data implicating a maladaptive role for alcohol in the neuropathological and behavioral developmental trajectory of AD. The 3xTg-AD mouse, and other models of AD, provide a unique opportunity to investigate how alcohol, and other modifiable risk factors, interact with mechanisms of AD and have potential to lead to synergistic discoveries that span research fields focused on age-dependent and drug-induced neurodegeneration. A potential future direction from these studies is to address mechanistic regulation by specific proteins, brain regions, and/or neural circuits. This could be accomplished by site-specific pharmacological, AAV, and optogenetic strategies. One circuit that might be important to evaluate is the projection from the lateral entorhinal cortex to the CA1 region of the HPC. The CA1 subregion shows intrinsic and circuit heterogeneity that may underlie the involvement of this structure in AD-associated behavioral pathologies (Masurkar, 2018); thus, subregional and circuit analyses have potential to identify specific anatomical substrates of alcohol’s impact on AD. Given the involvement of these brain regions in the onset of AD-like pathology, attenuating activity of this pathway may block or enhance disease state. Future studies might also pursue alcohol-induced instability of the Alzheimer’s-linked neural proteome or assess other mouse models of the disease. Additional approaches such as rodent fMRI to assess functional connectivity among AD target brain regions may also reveal novel neural pathways and mechanisms that are targeted by alcohol in AD-vulnerable or AD-expressing populations. Mechanistic studies are highly difficult, or even improbable, to conduct in human subjects; thus, further underscoring the value of having a well-characterized mouse model of AD to evaluate the impact of alcohol use on disease progression. Overall, mouse models of AD offer a novel and an exciting approach to assess the impact of alcohol use on neurodegeneration and represent an opportunity to identify novel therapeutic targets that could have long-lasting beneficial health consequences for the elderly population.

## AKNOWLEDGEMENTS

This research was supported by NIAAA grants R37AA014983, R37AA014983S1 and P60AA011065 to CW and by the UNC Bowles Center for Alcohol Studies. Mouse behavioral tests were conducted in the UNC Mouse Behavioral Core with assistance from Dr. Sheryl Moy supported by NICHD grant U54 HD079124. All authors declare that they have no conflict of interest.

## Notes

#### Summary of Updates

Clarification of methods, discussion of mouse models of Alzheimer's disease, and other editorial corrections.

